# Differentiation signals from glia are fine-tuned to set neuronal numbers during development

**DOI:** 10.1101/2021.12.13.472383

**Authors:** Anadika R. Prasad, Inês Lago-Baldaia, Matthew P. Bostock, Zaynab Housseini, Vilaiwan M. Fernandes

## Abstract

Neural circuit formation and function require that diverse neurons are specified in appropriate numbers. Known strategies for controlling neuronal numbers involve regulating either cell proliferation or survival. We used the *Drosophila* visual system to probe how neuronal numbers are set. Photoreceptors from the eye-disc induce their target field, the lamina, such that for every unit eye there is a corresponding lamina unit (column). Although each column initially contains ∼6 post-mitotic lamina precursors, only 5 differentiate into neurons, called L1-L5; the ‘extra’ precursor, which is invariantly positioned above the L5 neuron in each column, undergoes apoptosis. Here, we showed that a glial population called the outer chiasm giant glia (xg^O^), which resides below the lamina, secretes multiple ligands to induce L5 differentiation in response to EGF from photoreceptors. By forcing neuronal differentiation in the lamina, we uncovered that though fated to die, the ‘extra’ precursor is specified as an L5. Therefore, two precursors are specified as L5s but only one differentiates during normal development. We found that the row of precursors nearest to xg^O^ differentiate into L5s and, in turn, antagonise differentiation signalling to prevent the ‘extra’ precursors from differentiating, resulting in their death. Thus, an intricate interplay of glial signals and feedback from differentiating neurons defines an invariant and stereotyped pattern of neuronal differentiation and programmed cell death to ensure that lamina columns each contain exactly one L5 neuron.

## Introduction

Many sensory systems consist of repeated circuit units that map stimuli from the outside world onto sequential processing layers (Luo and Flanagan, 2007). It is critical that both absolute and relative neuronal numbers are carefully controlled for these circuits to assemble with topographic correspondence across processing layers. Neuronal numbers can be set by controlling how many progeny a neural stem cell produces, or by regulating how many neural progeny survive (Hidalgo, 2003; Miguel-Aliaga and Thor, 2009). To investigate other developmental strategies that set neuronal numbers, we used the highly ordered and repetitive *Drosophila melanogaster* visual system. Like vertebrate visual systems, the fly visual system is organized retinotopically into repeated modular circuits that process sensory input from unique points in space spanning the entire visual field (Hadjieconomou et al., 2011; Malin and Desplan, 2021).

Retinotopy between the compound eye and the first neuropil in the optic lobe, the lamina, is built during development. Photoreceptors are born progressively in the eye imaginal disc as a wave of differentiation sweeps across the tissue from posterior to anterior. Newly born photoreceptors express Hedgehog (Hh), which promotes further wave propagation (Treisman, 2013). They also express the Epidermal Growth Factor (EGF), Spitz (Spi), which recruits additional photoreceptors into developing ommatidia (Treisman, 2013). As photoreceptors are born, their axons project into the optic lobe and induce the lamina, such that there is a corresponding lamina unit (or cartridge) for every ommatidium (Figure 1A) (Hadjieconomou et al., 2011). Each cartridge is composed of five interneurons (L1-L5; named for the medulla layers they project to) and multiple glial subtypes (Fischbach and Dittrich, 1989; Hadjieconomou et al., 2011).

**Figure 1:**
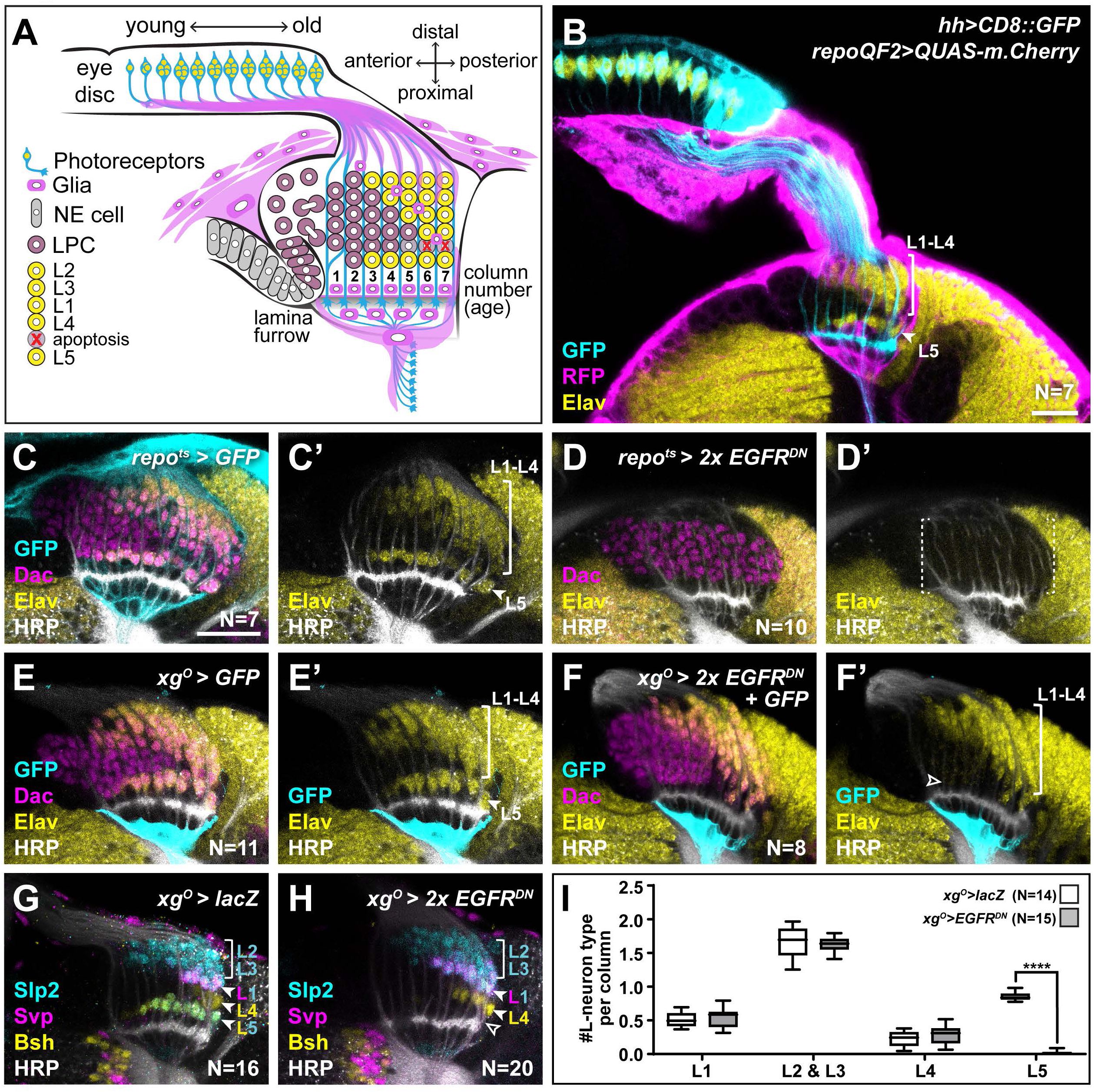
EGFR activity in the xg^O^ is required for the differentiation of L5 neurons. **(A)** Schematic of the developing lamina. Photoreceptors (blue) drive lamina precursor cell (LPC; purple) birth from neuroepithelial cells (NE; grey) and their assembly into columns of ∼6 LPCs, which differentiate into the L1-L5 neurons (yellow) following an invariant spatiotemporal pattern. The ‘extra’ LPC is cleared by apoptosis (red X). Several glial types (magenta) associate with the lamina. **(B)** A cross-sectional view of an early pupal (0-5 hours after puparium formation; APF) optic lobe where *hh-Gal4* drives *UAS-CD8::GFP* expression in photoreceptors (cyan). The pan-glial driver *repo-QF2* drives *QUAS-m.Cherry* (magenta) in all glia. Elav (yellow) marks all neurons. **(C)** A cross-sectional view of an optic lobe with pan-glial expression of CD8::GFP stained for GFP (cyan), Dac (magenta), Elav (yellow) and Horseradish peroxidase (HRP; axons; white). **(D)** Pan-glial expression of 2 copies of EGFR^DN^ stained for Dac (magenta), Elav (yellow) and HRP (white). (**E)** xg^O^-specific expression of CD8::GFP stained for GFP (cyan), Dac (magenta), Elav (yellow) and HRP (white). **(F)** xg^O^-specific expression of 2 copies of EGFR^DN^ and CD8::GFP stained for GFP (cyan), Dac (magenta), Elav (yellow) and HRP (white). The number of Elav+ cells in proximal row (L5s) decreased (empty arrowhead) relative to control (E). **(G,H)** HRP (white) and L-neuron type specific markers Slp2 (cyan), Bsh (yellow) and Svp (magenta) in **(G)** Control *xg^O^>lacZ* optic lobe and **(H)** *xg^O^>2xEGFR^DN^.* L2s and L3s express Slp2; L1s express Slp2 and Svp; L4s express Bsh and L5s express Bsh and Slp2. **(I)** Quantification of the number of L-neuron types per column for control and *xg^O^>2xEGFR^DN^.* Only L5 neurons were decreased significantly (P^L5^<0.0001; Mann-Whitney U test. Ns indicated in parentheses. Boxes indicate the lower and upper quartiles; the whiskers represent the minimum and maximum values; the line inside the box indicates the median). Scale Bar = 20µm.

Lamina induction is a multi-step process triggered by photoreceptor-derived signals. Photoreceptor-derived Hh converts neuroepithelial cells into lamina precursor cells (LPCs), promotes their terminal divisions and drives the assembly of lamina pre-cartridges referred to as columns, *i.e.* ensembles of ∼6 post-mitotic LPCs stacked together and associated with photoreceptor axon bundles (Figure 1A,B) (Huang and Kunes, 1998, 1996; Sugie et al., 2010; Umetsu et al., 2006). Once assembled into columns, LPCs are diversified by graded Hh signalling along the distal-proximal axis of young columns (Bostock et al., 2022). They then differentiate into neurons following an invariant spatio-temporal pattern whereby the most proximal (bottom) and most distal (top) cells differentiate first into L5 and L2, respectively; differentiation then proceeds in a distal-to-proximal (top-to-bottom) sequence, L3 forming next, followed by L1, then L4 (Fernandes et al., 2017; Huang et al., 1998; Tan et al., 2015). The sixth LPC, located between L4 and L5, does not differentiate but instead is fated to die by apoptosis and is later cleared (Figure 1A) (Apitz and Salecker, 2014). This spatio-temporal pattern of neuronal differentiation is driven in part by a population of glia called wrapping glia, which ensheathes photoreceptor axons and which induces L1-L4 neuronal differentiation via Insulin/ Insulin-like growth factor signalling in response to EGF from photoreceptors (Fernandes et al., 2017). Intriguingly, L1-L4 neuronal differentiation can be disrupted by manipulating wrapping glia without affecting L5 differentiation (Fernandes et al., 2017). Indeed, the mechanisms that drive L5 differentiation are not known. Importantly, we do not understand how exactly 5 neuron types differentiate from 6 LPCs; in other words, how are lamina neuronal numbers set?

Here, we sought to determine the mechanisms that drive L5 differentiation as well as those that set neuronal numbers in the lamina. We found that a population of glia located proximal to the lamina, called the outer chiasm giant glia (xg^O^), induces L5 neuronal differentiation in response to EGF from photoreceptors. We showed that the xg^O^ secrete multiple signals, including the EGF Spi and a type IV collagen, Collagen type IV alpha 1 (Col4a1), which activate Mitogen-Activated Protein Kinase (MAPK) signalling in the most proximal row of LPCs (*i.e.,* the row of LPCs nearest to xg^O^), thus driving their differentiation into L5s and promoting their survival. Further, we found that the ‘extra’ LPCs normally fated to die are specified with L5, but not L1-L4, identity. Since the most proximal row of LPCs are in closest proximity to the xg^O^, they receive differentiation cues from xg^O^ first and differentiate into L5s. In turn, these newly induced L5s secrete high levels of Argos (Aos), an antagonist of Spi (Freeman et al., 1992), to limit MAPK activity in the ‘extra’ LPCs thus preventing their differentiation, and leading to their death and clearance. Thus, we highlight a new mode by which neuronal numbers can be set – not only by regulating the number of neurons born or the number that survive, but also by regulating the number induced to differentiate from a larger pool of precursors. Altogether, our results indicate that the sterotyped pattern of neuronal differentiation and programmed cell death in the lamina are determined by the architecture of the developing tissue together with feedback from newly differentiating neurons.

## Results

### L5 neuronal differentiation requires EGF Receptor activity in outer chiasm giant glia (xg^O^)

We showed previously that wrapping glia induce L1-L4 neuronal differentiation in response to EGF from photoreceptors, but that L5 differentiation was regulated independently by an unknown mechanism (Fernandes et al., 2017). We speculated that another glial population may be involved in inducing L5 differentiation in response to EGF from photoreceptors. To test this hypothesis, we blocked EGF Receptor signalling in all glia using a pan-glial driver to express a dominant negative form of EGFR *(Repo>EGFR^DN^)*. Although LPCs (Dac+ cells) still formed and assembled into columns, there was a complete block in lamina neuron differentiation as seen by the absence of the pan-neuronal marker, Embryonic lethal abnormal vision (Elav); *i.e.,* L5 differentiation was disrupted in addition to the differentiation of L1-L4 as expected (Figure 1C, D). Thus, EGFR activity in a glial population other than the wrapping glia is required for L5 neuronal differentiation.

Many glial types infiltrate the lamina (Figure1 - Figure Supplement 1A) (Chotard and Salecker, 2007; Edwards et al., 2012). Therefore, we performed a screen using glia subtype-specific Gal4s to block EGFR signalling and determined what effect this manipulation had on L5s using Elav expression in the proximal lamina (Figure1 - Figure Supplement 1B-M; summarised in Supporting File 1). Blocking EGFR signalling in the outer chiasm giant glia (xg^O^) led to a dramatic reduction in the number of L5s (Figure 1E,F). To rule out early developmental defects, we used a temperature sensitive Gal80 (Gal80^ts^) and shifted animals from the permissive temperature to the restrictive temperature to limit EGFR^DN^ expression in xg^O^ to begin from the 3rd larval instar, when lamina development initiates. This resulted in a similar loss of Elav- positive cells in the proximal lamina as when EGFR^DN^ was expressed continuously in the xg^O^, indicating that this phenotype is not due to an early defect in xg^O^ (Figure 1 - Figure Supplement 1N). Xg^O^ are located below the lamina plexus, often with just one or two glial cells spanning the entire width of the lamina. While xg^O^ extend fine processes towards the lamina, they do not appear to contact LPCs or L5 neurons (Figure 1 - Figure Supplement 1O). Importantly, blocking EGFR signalling in the xg^O^ did not affect xg^O^ numbers or morphology (Figure 1E,F, Figure 1 - Figure Supplement 1O-R).

Since our screen used Elav expression in the proximal lamina to assess for the presence of L5s, we next examined lamina neuron type markers to assess whether blocking EGFR activity in xg^O^ affected L5 neurons specifically. We used antibodies against Sloppy paired 2 (Slp2), Brain-specific homeobox (Bsh) and Seven-up (Svp) in combination to distinguish lamina neuron types: L2s and L3s express Slp2 alone, L1s co-express Svp and Slp2, L4s express Bsh alone and L5s co-express Bsh and Slp2 (Figure 1G) (Fernandes et al., 2017; Hasegawa et al., 2013; Tan et al., 2015). We found that the number of L5 neurons decreased specifically, while the number of all the other neuron types were unaffected (Figure 1G-I; P^L5^<0.0001, Mann-Whitney U-test). Finally, to test whether the absence of L5s simply reflected a developmental delay in differentiation, we examined adult optic lobes using a different L5 neuronal marker, POU domain motif 3 (Pdm3) (Tan et al., 2015). Similar to our results in the developing lamina, L5s were mostly absent in the adult lamina when EGFR was blocked in xg^O^ compared with controls (Figure 1- Figure Supplement 1S,T; N^exp^=10; N^ctrl^=11), indicating that the loss of L5s observed during development is not due to delayed induction. Thus, EGFR activity in xg^O^ is required for L5 neuronal differentiation.

### LPCs that fail to differentiate as L5s are eliminated by apoptosis

The loss of L5 neurons when EGFR was blocked in xg^O^ could be explained either by a defect in neuronal differentiation, or by an earlier defect in LPC formation or recruitment to columns. To distinguish between these possibilities, we counted the number of LPCs per column when EGFR signalling was blocked in xg^O^ compared to controls (Figure 2A-C). For these and later analyses we considered the youngest column located adjacent to the lamina furrow to be the first column, with column number (and age) increasing towards the posterior side (Figure 1A). In columns 1-4, there were no differences in the number of LPCs when EGFR was blocked in xg^O^, indicating that LPC formation and column assembly occurred normally (Figure 2C), supporting the hypothesis that in response to EGFR activity, xg^O^ induce proximal LPCs to differentiate as L5s. Interestingly, the number of LPCs began to decrease in older columns (column 5 onwards) when EGFR signalling was blocked in xg^O^ (Figure 2C; P*<0.05, P****<0.0002, Mann-Whitney U-Test). This observation suggested that undifferentiated LPCs in older columns were being eliminated. We wondered whether LPCs that failed to differentiate into L5s underwent apoptosis, similar to the ‘extra’ LPCs that undergo apoptosis in controls. We used an antibody against the cleaved form of Death Caspase-1 (Dcp-1), an effector caspase, to detect apoptotic cells (Akagawa et al., 2015) and, indeed, observed a significant increase in the number of Dcp-1 positive cells in the lamina when EGFR signalling was blocked in the xg^O^ (132.8 cells/unit volume ± 19.48 standard error of the mean) compared to controls (49.14 cells/unit volume ± 4.53) (Figure 2A-B, 2D, P<0.0005, Mann-Whitney U Test). Importantly, we observed Dcp-1 positive cells in the proximal row of the lamina (Figure 2B; N^exp^=20/20), which we never observed in controls (Figure 2A, N^ctrl^=19/19). Altogether these results showed that EGFR activity in xg^O^ induces the differentiation of L5 neurons, and proximal LPCs that fail to receive appropriate cues from xg^O^ die by apoptosis.

**Figure 2:**
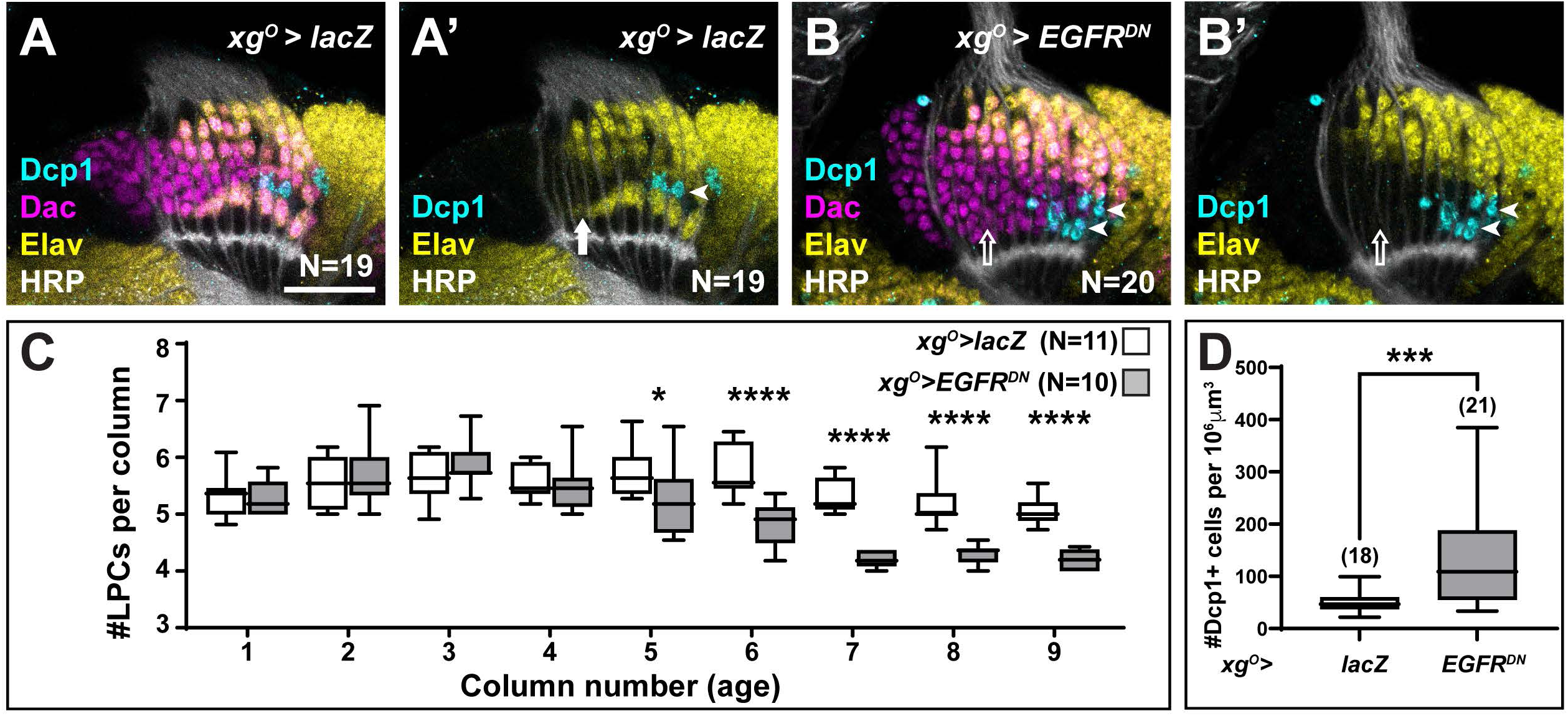
LPCs that fail to differentiate into L5s undergo apoptosis. **(A)** Control *xg^O^>lacZ* optic lobe stained for Dcp-1 (cyan), Elav (yellow) and HRP (white). Dcp- 1+ cells were always observed just distal to the most proximal row of cells (L5s). **(B)** *xg^O^>EGFR^DN^* stained for Dcp-1 (cyan), Dac (magenta) Elav (yellow) and HRP (white). Dcp-1 positive cells were observed in the most proximal row of LPCs as well as the row just distal to these. **(C)** Quantification of the number of LPCs/column (*i.e.* Dac+ cells/column) for control and *xg^O^>EGFR^DN^*. P*<0.05, P****<0.0002; Mann-Whitney U test. Ns indicated in parentheses. **(D)** Quantification of the number of Dcp-1 positive cells in (A) compared to (B). P***<0.0005, Mann-Whitney U test. Ns indicated in parentheses. Boxes indicate the lower and upper quartiles; the whiskers represent the minimum and maximum values; the line inside the box indicates the median). Scale bar = 20µm.

### xg^O^ respond to EGF from photoreceptors and secrete multiple ligands to induce MAPK-dependent neuronal differentiation of L5s

Since EGF from photoreceptors triggers EGFR activity in wrapping glia (Fernandes et al., 2017), we tested whether photoreceptor-derived EGF contributed to activating EGFR in xg^O^ also. Spi is initially produced as an inactive transmembrane precursor (mSpi) that needs to be cleaved into its active secreted form (sSpi) (Tsruya et al., 2002). This requires the intracellular trafficking protein Star and Rhomboid proteases (Tsruya et al., 2002; Urban et al., 2002; Yogev et al., 2008). We took advantage of a mutant for *rhomboid 3 (rho3)* in which photoreceptors are specified but cannot secrete EGF from their axons (Yogev et al., 2010), resulting in failure of L1-L4 neurons to differentiate along with a significant decrease in the number of L5s (Figure 3A, 3C; P*^rho3^*<0.0001; one-way ANOVA with Dunn’s multiple comparisons test) (Fernandes et al., 2017; Yogev et al., 2010). This result suggested that EGFR signalling in the xg^O^ could be activated by EGF secreted by photoreceptor axons. To test this hypothesis, we restored expression of wild-type Rho3 only in photoreceptors in *rho3* mutant animals using a photoreceptor-specific driver *(GMR-Gal4)*. Rho3 function in photoreceptors was sufficient to fully rescue not only L1-L4 neuronal differentiation, as previously reported (Yogev et al., 2010), but also L5 neuronal differentiation (Figure 3B, 3C; one-way ANOVA with Dunn’s multiple comparisons test). Since photoreceptor-derived EGF was insufficient to induce L5 neuronal differentiation when EGFR signalling was blocked in xg^O^ (Figure1 F,H), together these results suggest that xg^O^ likely respond to EGF from photoreceptors and relay these signals to induce differentiation of proximal LPCs into L5 neurons.

**Figure 3:**
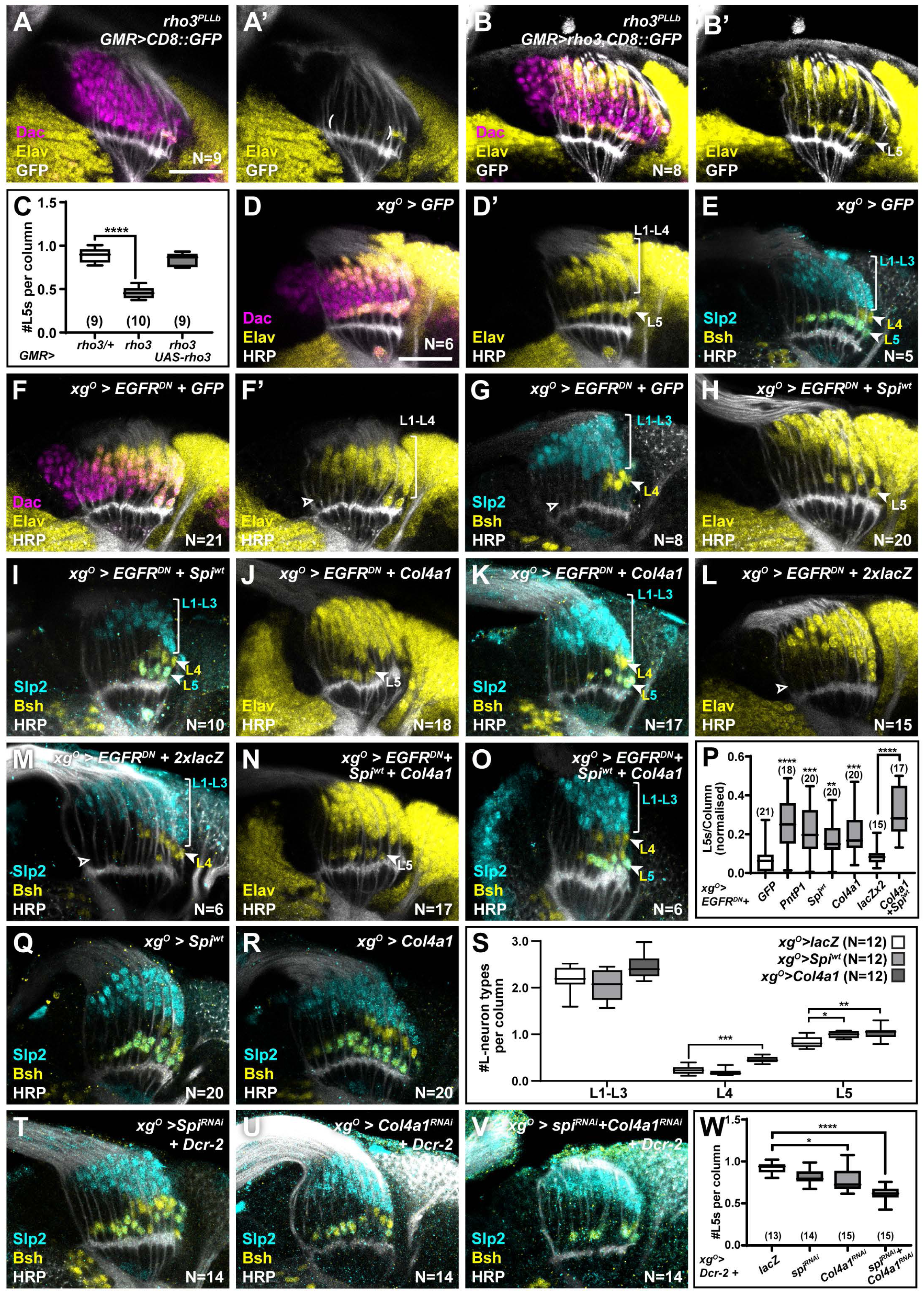
Xg^O^ secrete multiple ligands to induce L5 neuronal differentiation in response to EGF from photoreceptors. **(A)** GMR-Gal4 driven CD8::GFP expression in photoreceptors in a *rho3^PLLb^* background stained for GFP (white), Dac (magenta), Elav (yellow). Few proximal Elav+ cells (L5s) were recovered in older columns only as previously published (Fernandes et al., 2017). **(B)** GMR-Gal4 driven Rho3 and CD8::GFP in a *rho3^PLLb^* background stained for GFP (white), Dac (magenta), Elav (yellow) showed that L5 neuronal differentiation was rescued (Elav+ cells in the proximal lamina). **(C)** Quantifications for number of L5 neurons/column in (A) and (B) compared to *rho3^PLLb^* heterozygotes *(rho3*/+). P****<0.0001, one-way ANOVA with Dunn’s multiple comparisons test. Ns indicated in parentheses. **(D,E)** Control *xg^O^>GFP* optic lobes stained for **(D)** Dac (magenta), Elav (yellow) and HRP (white) or **(E)** HRP (white) and L-neuron specific markers Slp2 (cyan) and Bsh (yellow). **(F,G)** Gal4 titration control *xg^O^>GFP+EGFR^DN^* stained for **(F)** Dac (magenta), Elav (yellow) and HRP (white) or **(G)** HRP (white) and L-neuron specific markers Slp2 (cyan) and Bsh (yellow). **(H,I)** Wild-type Spi (Spi^wt^) co-expression with EGFR^DN^ specifically in xg^O^ stained for **(H)** Elav (yellow) and HRP (white) or **(I)** HRP (white) and L-neuron specific markers Slp2 (cyan) and Bsh (yellow). **(J,K)** Col4a1 co-expression with EGFR^DN^ specifically in xg^O^ stained for **(J)** Elav (yellow) and HRP (white) or **(K)** HRP (white) and L-neuron specific markers Slp2 (cyan) and Bsh (yellow). **(L,M)** Gal4 titration control *xg^O^>EGFR^DN^+2xlacZ* stained for **(L)** Elav (yellow) and HRP (white) or **(M)** HRP (white), Slp2 (cyan) and Bsh (yellow). **(N,O)** Wild-type Spi^wt^ and Col4a1 co-expression with EGFR^DN^ specifically in xg^O^. **(N)** stained for Elav (yellow) and HRP (white) or **(O)** HRP (white) and L-neuron specific markers Slp2 (cyan) and Bsh (yellow). **(P)** Quantification of the number of L5s/column for the genotypes indicated compared to the appropriate titration control. For *pntP1*, *spi^wt^*, and *Col4a1* co-expression with EGFR^DN^ the titration control is *xg^O^>EGFR^DN^+GFP* (P**<0.005, P***<0.0005; P****<0.0001; one-way ANOVA with Dunn’s multiple comparisons test. Ns indicated in parentheses). For *spi^wt^* and *Col4a1* simultaneous co-expression with EGFR^DN^ the titration control is *xg^O^>EGFR^DN^+2xLacZ* (P****<0.0001, Mann-Whitney U Test. Ns indicated in parentheses). **(Q,R)** Optic lobes stained for Slp2 and Bsh when xg^O^ overexpress **(Q)** *spi^wt^* or **(R)** *Col4a1*. **(S)** Quantification of the number of L-neuron types/column in (Q) and (R) compared to controls, *xg^O^>lacZ*. (P*<0.05; P**<0.005; P***<0.001; one-way ANOVA with multiple comparisons test). **(T, U, V)** Optic lobes stained for Slp2, Bsh and HRP when xg^O^ co-express Dcr-2 with **(T)** spi^RNAi^, **(U)** Col4a1^RNAi^ and **(V)** Spi^RNAi^ and Col4a1^RNAi^ simultaneously. **(W)** Quantifications of the number of L5s/column for genotypes indicated compared to the titration control *xg^O^>Dcr-2+lacZ* (P*<0.05, P****<0.0001, one-way ANOVA with Dunn’s multiple comparisons test. Scale bar = 20µm. For all quantifications boxes indicate the lower and upper quartiles; the whiskers represent the minimum and maximum values; the line inside the box indicates the median).

We next asked what signal(s) the xg^O^ secrete to induce L5 differentiation. Previously, we showed that MAPK signalling is necessary and sufficient for neuronal differentiation in the lamina (Fernandes et al., 2017). Therefore, we reasoned that xg^O^-derived differentiation signal(s) must activate MAPK signalling through a Receptor Tyrosine Kinase (RTK) in the proximal lamina. Indeed, blocking EGFR signalling in xg^O^ led to reduced levels of double phosphorylated MAPK (dpMAPK) specifically in the proximal lamina (Figure 3 - Figure Supplement 1A, B). The *Drosophila* genome encodes 22 ligands which activate 10 RTKs upstream of MAPK signalling (Sopko and Perrimon, 2013). To identify the signal(s) secreted by xg^O^, we misexpressed candidate ligands and screened for their ability to rescue the loss of L5s caused by blocking EGFR activity in the xg^O^. To validate this approach, we tested whether autonomously restoring transcriptional activity downstream of MAPK in xg^O^ while blocking EGFR activity could rescue L5 differentiation. While blocking EGFR in xg^O^ resulted in laminas containing 0.063 ± 0.014 L5s per column, co-expressing PntP1 with EGFR^DN^ in xg^O^ rescued the number of L5s per column to 0.213 ± 0.025 (Figure 3F, Figure 3 - Figure Supplement 1C P****<0.0001 compared to EGFR^DN^ alone). We then screened 18 RTK ligands based on available reagents (Figure 3 - Figure Supplement 1C). Four ligands, Spitz (Spi), Branchless (Bnl), Thisbe (Ths) and Collagen type IV alpha 1 (Col4a1) produced statistically significant rescues when compared with the *xg^O^>EGFR^DN^+CD8::GFP* (Gal4 titration control) (Figure 3- Figure Supplement 1C; P*< 0.05, P**<0.005, P***<0.0005, P****<0.0001 one-way ANOVA with Dunn’s multiple comparisons test). To eliminate false positive hits, we determined whether these ligands were expressed in xg^O^ under physiological conditions. Using a previously validated *bnl-Gal4* (Chen and Krasnow, 2014; Kamimura et al., 2006; Spéder and Brand, 2014; Tamamouna et al., 2021), we drove CD8::GFP expression and found that it was expressed in all cells of the optic lobe (Figure 3 - Figure Supplement 1D), making it unlikely to be a viable hit. We found that a previously validated *ths-Gal4* (Anllo and DiNardo, 2022; Wu et al., 2017) drove CD8::GFP expression in photoreceptors but not xg^O^ (Figure 3 - Figure Supplement 1E) consistent with previous reports (Franzdóttir et al., 2009). However, when we examined *Col4a1* expression using a previously validated Gal4 enhancer trap (Hennig et al., 2006), we found that it drove CD8::GFP expression in xg^O^ (Figure 3 - Figure Supplement 1F). We also found that a *spi-Gal4* (NP0289-Gal4; not previously validated) drove CD8::GFP expression in xg^O^, but not photoreceptors or other cell types where *spi* is also known to be expressed, suggesting that this Gal4 line may report *spi* expression partially (Figure 3 - Figure Supplement 1G). To further substantiate these results we performed fluorescence *in situ* hybridisation chain reaction (HCR), a form of fluorescent *in situ* hybridisation (Choi et al., 2018, 2016; Duckhorn et al., 2022), and confirmed that *spi* and *Col4a1* mRNAs were present in the xg^O^ under physiological conditions (Figure 3 - Figure Supplement 1H, 1I; See Materials and Methods). This enabled us to narrow down our hits to two ligands: the EGF Spi and Col4a1, a type IV collagen, which both rescued L5 differentiation resulting in laminas with 0.147 ± 0.024 and 0.17 ± 0.0197 L5s per column, respectively (Figure 3F-K, 3P P^spi-wt^<0.01 and P^Col4a1^ <0.0005, one-way ANOVA with Dunn’s multiple comparisons test; Figure 3- Figure Supplement 1C). Note that expressing either sSpi or wild-type (unprocessed) mSpi (referred to as Spi^wt^) in xg^O^ rescued L5 numbers (Figure 3 – Figure supplement 1C), indicating that xg^O^ are capable of processing mSpi into the active form (sSpi).

We ruled out the trivial explanation that the rescue of L5 numbers by Spi was caused by autocrine EGFR reactivation in the xg^O^, as Spi expression in xg^O^ did not autonomously rescue dpMAPK nuclear localisation when EGFR signalling was blocked (Figure 3 – Figure Supplement 1A, B, J, K). We then tested whether xg^O^ express *spi* and *Col4a1* downstream of EGFR activity. We measured *spi* and *Col4a1* transcript levels using *in situ* HCR in controls and when we blocked EGFR signalling in xg^O^. Disrupting EGFR signalling in xg^O^ resulted in a significantly reduced fluorescence signal for *spi* and *Col4a1* transcripts in xg^O^ compared with controls (Figure 3- Figure Supplement 1H, 1I,1L-O; P*^spi^*<0.01, P*^Col4a1^*<0.005; Mann-Whitney U Test). Thus, xg^O^ express *spi* and *Col4a1* in response to EGFR activity.

Col4a1 is thought to activate MAPK signalling through its putative receptor, the Discoidin domain receptor (Ddr) (Sopko and Perrimon, 2013). We used a Gal4 enhancer trap in the *Ddr* locus (not previously validated) to drive CD8::GFP expression and observed that GFP was expressed in all LPCs (Figure 3- Figure Supplement 1P). We confirmed these results using *in situ* HCR, which also detected *Ddr* expression throughout the lamina (Figure 3 – Figure supplement 1Q). Spi activates EGFR (Sopko and Perrimon, 2013), which was shown to be expressed in LPCs previously (Huang et al., 1998). Thus, LPCs express the RTKs that make them competent to respond to the EGF Spi and Col4a1 produced by xg^O^. Moreover, expressing *spi* or *Col4a1* in xg^O^ in which EGFR signalling was blocked rescued dpMAPK signal in L5s, indicating that, when expressed in xg^O^, these ligands were sufficient to activate MAPK signalling in the proximal lamina (Figure3-Figure Supplement R-T; P**<0.005, P****<0.0001; one-way ANOVA with Dunn’s multiple comparisons Test). Co-expressing Spi and Col4a1 in the *xg^O^>EGFR^DN^* background led to an enhanced and statistically significant rescue relative to individual ligand rescues alone, resulting in laminas with 0.267 ± 0.025 L5s per column (Figure 3L-P; P<0.0001, Mann-Whitney U Test). We also tested whether these ligands could induce ectopic L5 differentiation when overexpressed in the xg^O^. Overexpressing either Spi or Col4a1 resulted in a 19% ± 1.8 (P<0.05) and a 24% ± 4 (P<0.005) increase in the number of L5s per column relative to controls, respectively (Figure 3Q-S). Thus, Spi and Col4a1 from xg^O^ are sufficient to induce L5 differentiation.

Next, to test whether xg^O^-derived Spi and Col4a1 are normally required to induce L5 neuronal differentiation, we disrupted their expression specifically in xg^O^. We used RNA interference (RNAi) to knock down *spi* and *Col4a1* expression both individually and simultaneously in xg^O^ using previously validated lines (Chen et al., 2016; Csordás et al., 2020; Louradour et al., 2017; Morante et al., 2013; Pastor-Pareja and Xu, 2011). While knocking down *spi* led to a mild decrease in L5 numbers, which was not statistically significant, knocking down *Col4a1* in the xg^O^ led to a statistically significant decrease in L5s (0.78 ± 0.03 L5s per column) relative to controls (0.92 ± 0.02 L5s per column) (Figure 3T, 3U, 3W; P*<0.05 one-way ANOVA with Dunn’s multiple comparisons test). However, knocking down both *spi* and *Col4a1* simultaneously in xg^O^ led to a strong decrease in L5s (0.61 ± 0.02 L5s per column; Figure 3V- W; P****<0.0001,one-way ANOVA with Dunn’s multiple comparisons test). Under these conditions we also observed Dcp1 positive apoptotic cells in the most proximal row of the lamina, which were never observed in controls (Figure 3- Figure supplement 2) but were observed when L5 differentiation was blocked above (Figure 2B). Thus, xg^O^-derived Spi and Col4a1 are both necessary and sufficient to induce L5 differentiation. Altogether, we found that xg^O^ secrete multiple factors that lead to activation of the MAPK cascade in the proximal lamina to induce differentiation of L5s.

### The ‘extra’ lamina precursor cells are specified as L5s though fated to die

We recently showed that a gradient of Hh signalling activity in lamina columns specifies L1-L5 identities such that high levels specify L2 and L3 (distal cell) identities, intermediate levels specify L1 and L4 (intermediate cell) identities and low levels specify L5 (proximal cell) identity (Bostock et al., 2022). Since overexpressing *spi* and *Col4a1* in the xg^O^ resulted in ectopic L5 neurons, we wondered what the source of these ectopic cells was. We quantified other lamina neuron types when either *spi* or *Col4a1* was overexpressed in xg^O^ and found no decrease in the number of L1-L3s (Slp2-only expressing cells) or L4s (Bsh-only expressing cells) per column compared to controls (Figure 3S). Thus, ectopic L5s were not produced at the expense of other lamina neuron types. In wild-type optic lobes, each lamina column contains an ‘extra’ LPC, which is located immediately distal to the LPC fated to differentiate as an L5. These ‘extra’ LPCs do not differentiate but instead undergo apoptosis and are eliminated (Figure 2A and 4A). We hypothesised that though fated to die, ‘extra’ LPCs are specified with L5 identity through low Hh signalling activity in the proximal lamina, and that the overexpression of Spi and Col4a1 in xg^O^ generated ectopic L5s by inducing differentiation and survival of the ‘extra’ LPCs. To test this hypothesis, we forced neuronal differentiation throughout the lamina by expressing an activated form of MAPK (MAPK^ACT^) (Figure 4 – Figure Supplement 1A-D) or by overexpressing the MAPK transcriptional effector, Pointed P1 (PntP1), in the lamina (Figure 4A- D). As reported previously, hyperactivating MAPK signalling in the lamina led to premature neuronal differentiation: instead of sequential differentiation of L1-L4, seen as a triangular front, most lamina columns differentiated simultaneously (Figure 4C, Figure 4 - Figure Supplement 1A, C) (Fernandes et al., 2017). We observed no LPCs that remained undifferentiated (Dac^+^ and Elav^-^) past lamina column 5, including the row of cells that normally correspond to the ‘extra’ LPCs (Figure 4C, Figure 4 - Figure Supplement 1A, C). Importantly, we also observed a concomitant decrease in cleaved Dcp-1 positive cells (Figure 4C, E ; P<0.0001, Mann-Whitney U Test), suggesting that forcing the ‘extra’ LPCs to differentiate blocked their death.

**Figure 4:**
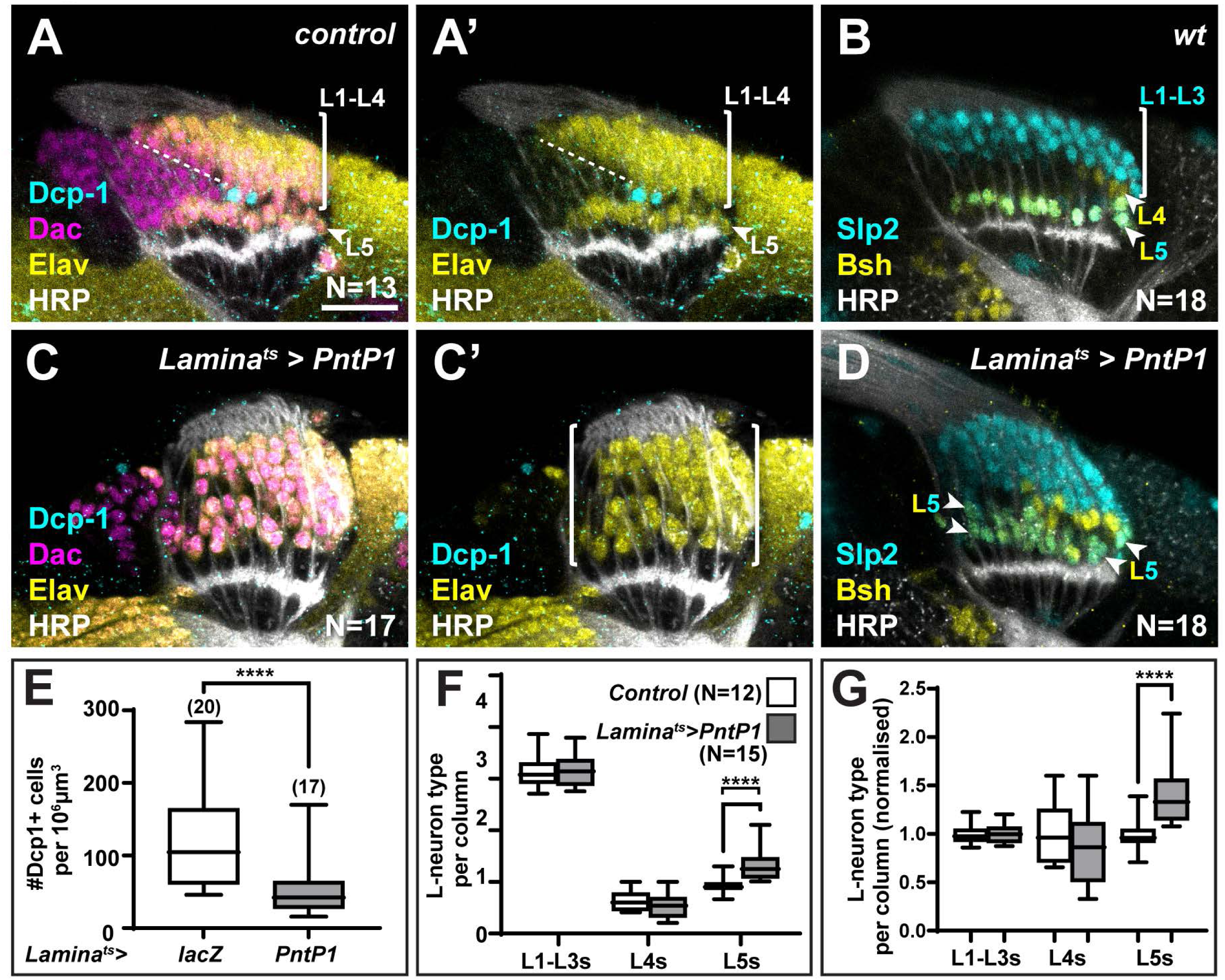
The ‘extra’ LPCs are specified as L5s. **(A)** Wild-type optic lobes stained for Dac (magenta), HRP (white), Elav (yellow) and cleaved Dcp-1 (cyan). **(B)** Wild-type optic lobes stained for HRP (white) and L-neuron type specific markers Slp2 (cyan) and Bsh (yellow). **(C, D)** Optic lobes with lamina-specific overexpression of PntP1 stained as in (A) and (B), respectively. **(C)** Fewer Dcp-1 positive cells were recovered compared with controls. **(D)** Roughly two rows of Slp2 and Bsh co-expressing cells (L5s) were recovered (arrow heads). **(E)** Quantification of the number of Dcp-1 positive cells in (B) compared with control *Lamina^ts^>lacZ* (Figure 4- figure supplement 1A) (P<0.0001; Mann-Whitney U test). **(F)** Quantification of the number of L-neuron types per column based on Slp2 and Bsh expression from column 7 onwards shows an increase in the number of L5s/column in *Lamina^ts^>PntP1* compared with controls; P<0.0001; Mann-Whitney U test. **(G)** Same as (F) but normalised to the mean of the control. The number of L5s/column in *Lamina^ts^>PntP1* increase ∼1.2 fold relative to controls; P <0.0001; Mann-Whitney U Test. Ns indicated in parentheses. Scale bar = 20µm. For all quantifications boxes indicate the lower and upper quartiles; the whiskers represent the minimum and maximum values; the line inside the box indicates the median).

Next, we examined the distribution of lamina neuron types when we forced neuronal differentiation. We often observed two rows of cells co-expressing Slp2 and Bsh in the proximal lamina (Figure 4B, D, Figure 4- figure supplement 1B, D), indicating the presence of ectopic L5s. To distinguish between premature and ectopic differentiation, we quantified the number of lamina neuron types (L1-L3, L4 and L5) per column in older columns (column 7 onwards, once mature columns were observed in controls, Figure 4 - Figure Supplement 1E). While there was no significant difference between the average number of L1-L3s or L4s per column, the average number of L5s per column was ∼1.4-fold higher in laminas in which differentiation was ectopically induced compared with controls, *i.e.* they contained 1.4 ± 0.08 L5s per column compared to 1.00 ± 0.05 L5s per column in controls (Figure 4B, D, F-G; P<0.0001, Mann- Whitney U Test). Thus, hyperactivating MAPK signalling in the lamina drove ectopic differentiation of L5 neurons. Importantly, ectopic L5s were only observed in the proximal but never in the distal lamina (Figure 4D, N=18/18; Figure 4 - Figure Supplement 1D, N=9/9).

Taken together, the absence of cell death in the row distal to L5s and the presence of ectopic L5s in this row indicate that hyperactivating MAPK signalling induces the ‘extra’ LPCs to differentiate into L5s. Thus, the ‘extra’ LPCs are specified as L5s though fated to die normally. These data are consistent with our work showing that lamina precursors are specified by Hh signalling prior to differentiation and that the most proximal cells, which experience the lowest levels of Hh pathway activity and are specified as L5s (Bostock et al., 2022). Importantly, the presence of ectopic L5s when differentiation is induced demonstrates that more LPCs are specified as L5s than differentiate normally.

### Newly born L5 neurons inhibit differentiation of distal neighbours to set neuronal number

If the two most proximal cells in each lamina column are both specified as L5s, how then is L5 differentiation limited to only the most proximal row in response to diffusible signals secreted by xg^O^? We tested whether the ‘extra’ LPCs differentiated as L5s when apoptosis was blocked in animals mutant for *Death regulator Nedd2-like caspase (Dronc)*, an initiator caspase essential for caspase-dependent cell death (Fuchs and Steller, 2011). Cleaved Dcp-1 was absent in homozygous *Dronc^I24^* animals confirming that apoptosis was blocked (Figure 5A; N=26/26; with full penetrance). Indeed, we detected cells that were positive for the lamina marker Dachshund (Dac) but negative for the pan-neuronal marker Elav between L1-L4 and L5 neurons past column 5, which were never observed in controls (Figure 5A compared to 4A; N=13/13; with full penetrance). These cells did not express lamina neuron type markers Slp2 or Bsh, which L5s co-express and which individually label L1-L3s and L4s, respectively (Figure 5B,C). Thus, although the ‘extra’ LPCs were retained when apoptosis was blocked, they did not differentiate into neurons.

**Figure 5:**
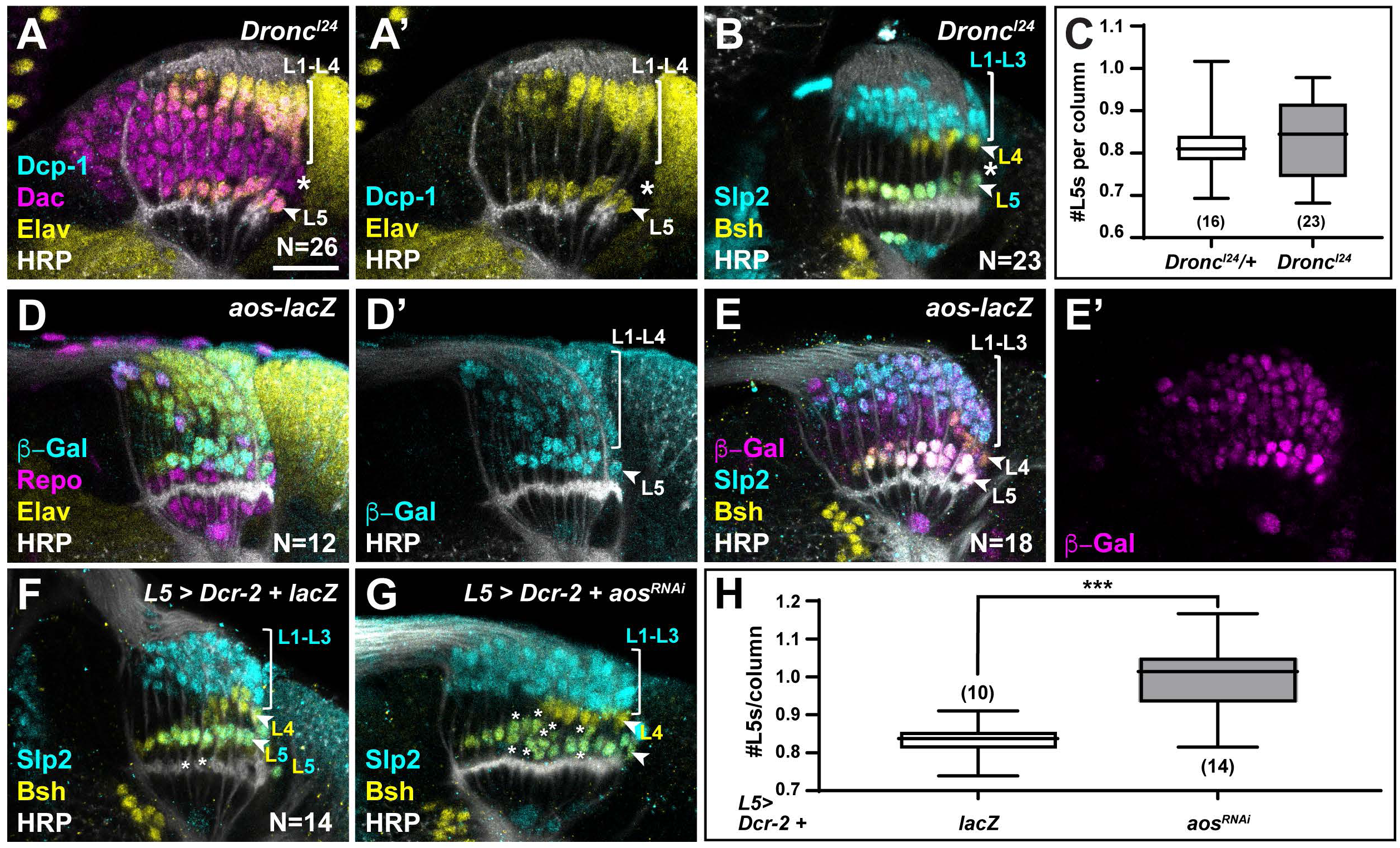
Newly-induced L5 neurons secrete Aos to limit differentiation signals from xg^O^. **(A)** *Dronc^I24^* optic lobes stained for Dcp-1 (cyan), Dac (magenta), Elav (yellow) and HRP (white). No Dcp-1 positive cells were recovered and Dac positive cells between L1-L4s and L5s persisted into the oldest columns (asterisk). **(B)** *Dronc^I24^* optic lobes stained for L-neuron type specific markers Slp2 (cyan) and Bsh (yellow). A space (negative for both markers; asterisk) was present between L4s and L5s. **(C)** Quantifications for number of L5s/column in *Dronc^I24^* optic lobes compared to controls (*Dronc^I24^/+*) (P>0.05, Mann-Whitney U test. Ns indicated in parentheses). **(D,E)** *aos-lacZ* expression in the lamina with **(D)** b-Gal (cyan), Repo (magenta), Elav (yellow), HRP (white) and with **(E)** b-Gal (magenta) and L-neuron type specific markers Slp2 (cyan), Bsh (yellow) as well as HRP (white). **(F)** An L5-specific Gal4 was used to drive expression of *Dcr-2* and *lacZ* in control lobes stained for Slp2 (cyan), Bsh (yellow) and HRP (white). **(G)** Optic lobes stained for HRP (white), Slp2 (cyan) and Bsh (yellow) when *Dcr-2* and *aos^RNAi^* were expressed in developing L5 neurons specifically, which led to an increase in the number of Slp2 and Bsh co-expressing cells (L5s; asterisks). **(H)** Quantification of the number of L5s/column for (F) and (G). P***<0.0005; Mann-Whitney U test. Ns indicated in parentheses. For all quantifications boxes indicate the lower and upper quartiles; the whiskers represent the minimum and maximum values; the line inside the box indicates the median). Scale bar = 20µm.

**Figure 6:**
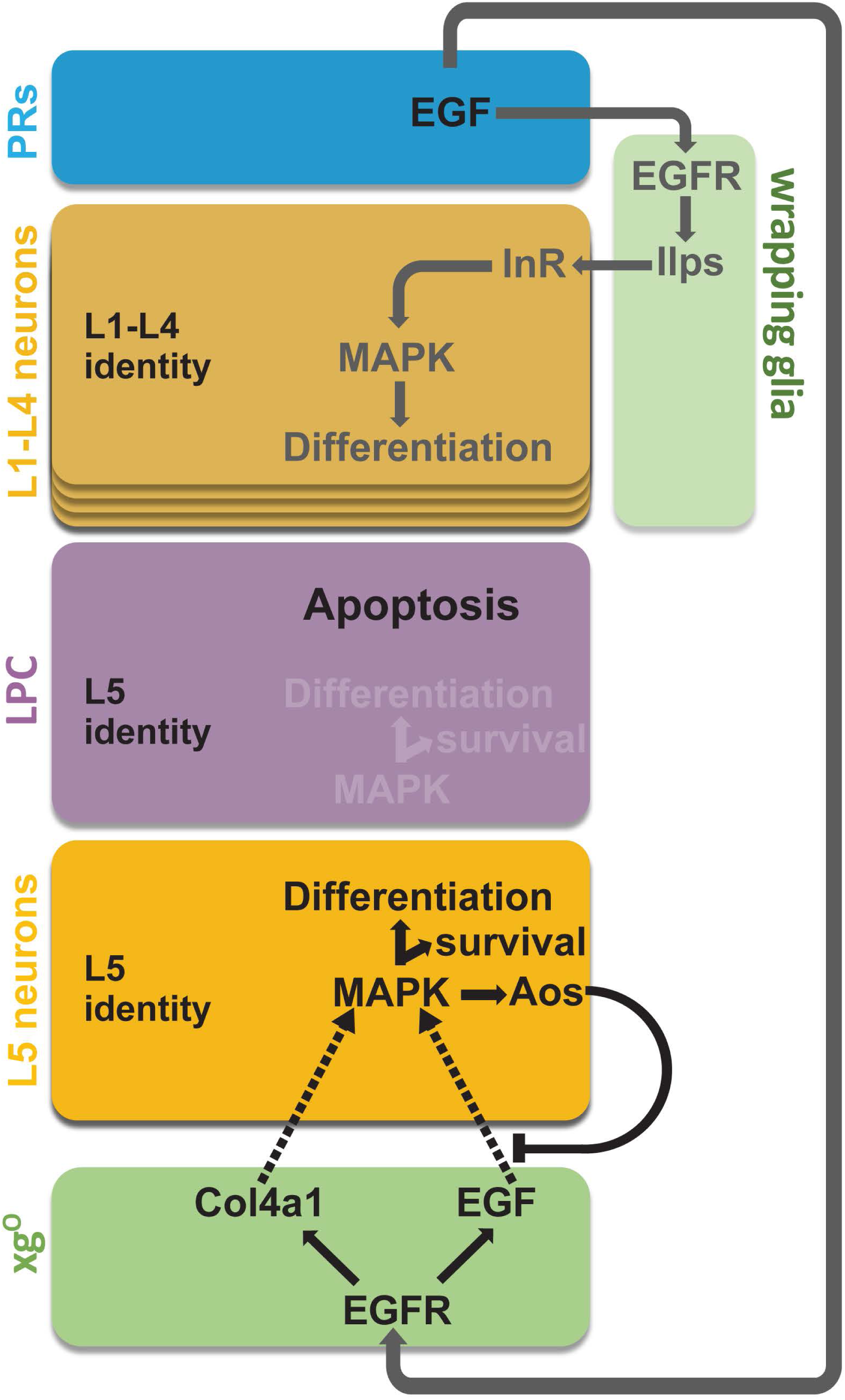
Summary schematic of neuronal differentiation in the lamina. In our model of lamina neuronal differentiation, lamina precursor cells are prepatterned with unique identities based on their positions within a column, such that the two most proximal cells are specified with L5 identity. EGF from photoreceptors activates EGFR signalling in wrapping glia, which induce L1-L4 differentiation, and in xg^O^, which induce L5 differentiation. Only a subset of the LPCs specified as L5s differentiate (*i.e.* those in the proximal row). We propose that this selective neuronal induction of L5s is due to tissue architecture and feedback from the newly born L5s, which limit available EGF (Spi) by secreting the antagonist Aos.

We observed ectopic L5s only when all LPCs were forced to differentiate, bypassing the need for differentiation signals from glia, but not when apoptosis was blocked (Figure 4C-D, Figure 4-figure supplement 1A-D and Figure 5A-C). This suggests that the ‘extra’ LPCs, though specified as L5s, did not receive differentiation signals from xg^O^ in *Dronc* mutants or in the wild-type, where failing to differentiate they were eliminated by apoptosis. How are only half of the LPCs specified as L5s chosen to differentiate in an invariant manner? The most proximal row of LPCs fated to differentiate into L5s is the row nearest to xg^O^, and therefore, the first to receive differentiation signals. We speculated that newly-induced L5s may then limit the ability of more distal LPCs to differentiate, by preventing MAPK activation in neighbouring cells. Aos is a transcriptional target of MAPK signalling and a secreted antagonist of the EGF Spi (Freeman et al., 1992; Golembo et al., 1996). We wondered if newly-induced L5s secrete Aos to limit differentiation signals from xg^O^. To test this hypothesis, we examined *argos* (*aos*) expression with an enhancer trap in the *aos* locus, *aos^W11^. aos-lacZ (aos^W11/+^)* was expressed in xg^O^ and differentiating lamina neurons, with the highest levels detected in L5s (Figure 5D-E). Interestingly, we also noted ectopic L5s in the laminas of *aos^W11^* heterozygotes, which could be the result of decreased Aos expression, as *aos^W11^* is a hypomorphic loss-of-function allele (Figure 5E). These observations suggested that Aos could act in L5s as a feedback-induced sink for Spi to limit further differentiation in columns. To test this hypothesis, we knocked down *aos* by RNAi using a driver expressed specifically in developing L5s (Jenett et al., 2012) (Figure 5G). We observed a statistically significant ∼1.2-fold increase in the number of L5s relative to controls, *i.e.* 0.99 ± 0.02 L5s per column compared to 0.83 ± 0.01 L5s per column in controls (Figure 5F-H; P<0.0005, Mann-Whitney U Test). Altogether, out data indicate a model in which xg^O^ induce MAPK activity in the most proximal LPCs, resulting in their differentiation and in the production of the feedback inhibitor Aos. In turn, Aos limits further differentiation in the column by fine-tuning the availability of the differentiation signal Spi, which ensures that only one L5 differentiates per column, and determines the final number of neurons in each lamina column.

## Discussion

Appropriate circuit formation and function require that neuronal numbers are tightly regulated. This is particularly important for the visual system, which is composed of repeated modular circuits spanning multiple processing layers. In *Drosophila,* photoreceptors induce their target field, the lamina, thus, establishing retinotopy between the compound eye and the lamina (Huang and Kunes, 1996). Each lamina unit or column in the adult is composed of exactly 5 neurons; however, columns initially contain 6 LPCs. The sixth, or ‘extra’, LPC, invariantly located immediately distal to the differentiating L5 neuron, is fated to die by apoptosis. These ‘extra’ LPCs did not differentiate when apoptosis was blocked (Figure 5A,B) but generated ectopic L5s when forced to differentiate (Figure 4D and Figure 4- figure supplement 1D). Although we cannot rule out that preventing death using *Dronc* mutants may mis-specify the ‘extra’ cells and prevent them from differentiating, it is more likely that these ‘extra’ cells are specified as L5s, but that other mechanisms restricted their differentiation in *Dronc* mutants, as other lamina neuron types differentiated normally (Figure 5B). Thus, twice as many LPCs appear to be specified as L5s than undergo differentiation normally, which implies that a selection process to ensure the correct number of L5s develop is in place.

The developmental strategies described thus far for setting neuronal number do so by regulating proliferation of precursors and/or survival of differentiated neurons (Hidalgo, 2003). Here, we have defined a unique strategy whereby L5 neuronal numbers are set by regulating how many precursors from a larger pool are induced to differentiate, followed by programmed cell death of the excess precursors. We showed that a glial population called xg^O^, which are located proximal to the lamina, secrete at least two ligands (Spi, Col4a1) that activate MAPK signalling in LPCs to induce their differentiation (Figure 3, Figure 3 - Figure Supplement 1). The tissue architecture is such that secreted signals from the xg^O^ reach the most proximal row of LPCs first, and therefore these precursors differentiate first. Upon differentiation, these newly induced neurons secrete the Spi antagonist Aos to limit the available pool of Spi. As a result, the MAPK pathway is not activated in the ‘extra’ L5 LPCs, preventing them from differentiating into L5 neurons (Figure 5). Intriguingly, L5 neuronal differentiation in the youngest columns of the lamina proceeds despite Aos secretion by newly-induced L5s. We noted that differentiating L5s expressed *aos* (based on *aos-lacZ*) at low levels initially and increased expression gradually till it plateaued from column 5 onwards (Figure 5E and Figure 5 – Figure Supplement 1B). This delay in high *aos* expression may thus enable differentiation of the youngest LPCs, while still inhibiting differentiation of the row immediately distal. In sum, the structure of the tissue together with feedback from newly induced neurons set neuronal number by limiting which and, therefore, how many LPCs are induced to differentiate.

### Co-ordinating development through glia

We have shown that in addition to the wrapping glia (Fernandes et al., 2017), another population of glia, the xg^O^, also receive and relay signals from photoreceptors to induce neuronal differentiation in the lamina (Figure 1E-F). This is the first functional role ascribed to xg^O^. Remarkably, xg^O^ are born from central brain DL1 type II neuroblasts and migrate into the optic lobes to positions below the developing lamina (Ren et al., 2018; Viktorin et al., 2013). This underscores an extraordinary degree of coordination and interdependence between the compound eye, optic lobe and central brain. Photoreceptor signals drive wrapping glial morphogenesis and infiltration into the lamina (Franzdóttir et al., 2009), thus setting the pace of L1-L4 neuronal differentiation (Fernandes et al., 2017). Defining the signals that enable xg^O^ to navigate the central brain and optic lobe will be a critical contribution to our understanding of how development is coordinated across brain regions.

### Tissue architecture sets up stereotyped programmed cell death

In both vertebrate and invertebrate developing nervous systems, programmed cell death is thought to come in two broad flavours: first as an intrinsically programmed fate whereby specific lineages or identifiable progenitors, neurons or glia undergo stereotyped clearance (Hidalgo, 2003; Miguel-Aliaga and Thor, 2009; Pinto-Teixeira et al., 2016; Yamaguchi and Miura, 2015) and second as an extrinsically controlled outcome of competition among neurons for limited target-derived trophic factors, which adjust overall cell numbers through stochastic clearance (also known as the neurotrophic theory) (Davies, 2003; Hidalgo, 2003; Miguel-Aliaga and Thor, 2009; Yamaguchi and Miura, 2015). In the lamina, although the LPCs eliminated by programmed cell death are identifiable and the process stereotyped, it does not appear to be linked to an intrinsic programme. Rather, the predictable and stereotyped nature of apoptosis and differentiation are a consequence of stereotyped responses to extrinsic signalling determined by the architecture of the tissue. Thus, our work highlights that stereotyped patterns of apoptosis can arise from extrinsic signalling, suggesting a new mode to reliably pattern development of the nervous system.

In many contexts, neurotrophic factors promote cell survival by activating MAPK signalling (Ballif and Blenis, 2001; Park and Poo, 2013). In the lamina, MAPK-induced neuronal differentiation and cell survival appear intimately linked. LPCs that do not activate MAPK signalling sufficiently do not differentiate and are eliminated by apoptosis, likely through regulation of the proapoptotic factor Head involution defective, which has been described extensively in flies (Bergmann et al., 2002, 1998; Kurada and White, 1998). Thus, here the xg^O^-secreted ligands Spi and Col4a1, which activate MAPK, appear to be functioning as differentiation signals as well as trophic factors. Col4a1, in particular, may perform dual roles by promoting MAPK activity directly through its receptor Ddr, and perhaps also by limiting Spi diffusivity to aid in localising MAPK activation.

It will be interesting to determine whether the processes described here represent conserved strategies for regulating neuronal number. Certainly, given the diversity of cell types and structural complexity of vertebrate nervous systems, exploiting tissue architecture would appear to be an effective and elegant strategy to regulate cell numbers reliably and precisely.

## Supporting information

Supporting zip file 1

## Acknowledgements

We thank C. Desplan, A. Gould, B. Shilo and J. Treisman for reagents, and S. Ackerman, M. Amoyel, B. Conradt, C. Desplan, C. Doe, A. Franz, P. Salinas, A. Rossi, C. Stern, L. Venkatasubramanian and members of the Amoyel and Fernandes labs for comments on the manuscript. Stocks obtained from the Bloomington Drosophila Stock Center (NIH P40OD018537) were used in this study. Monoclonal antibodies obtained from the Developmental Studies Hybridoma Bank, created by the NICHD of the NIH and maintained at The University of Iowa, were used in this study.

## Author contributions

Conceptualization: VMF, ARP Investigation: ARP, MPB, ILB, ZH, VMF Supervision: VMF Writing – original draft: VMF, ARP Writing – review & editing: ARP, ILB als and Methods

## Materials and Methods

### Key Resources Table

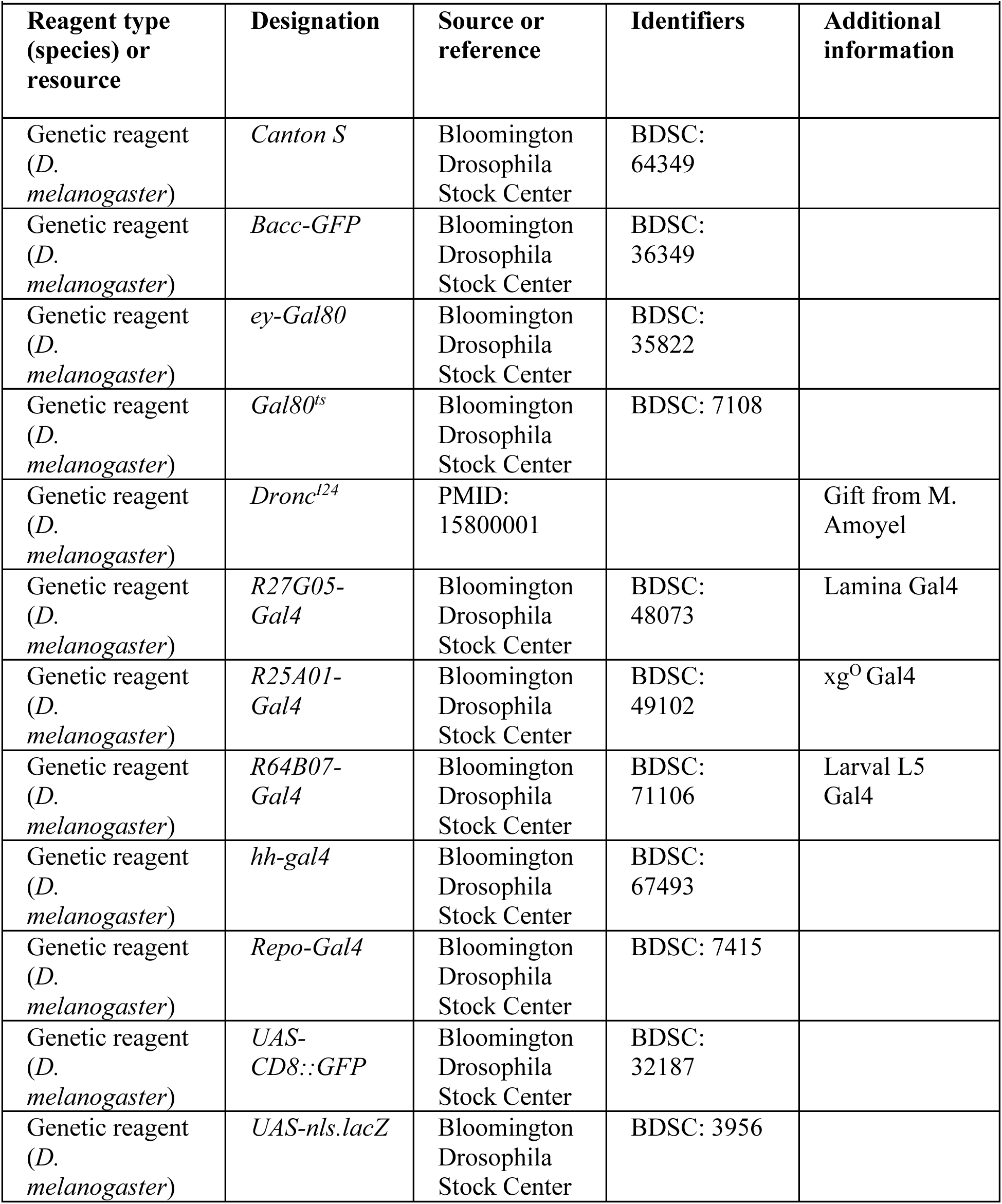
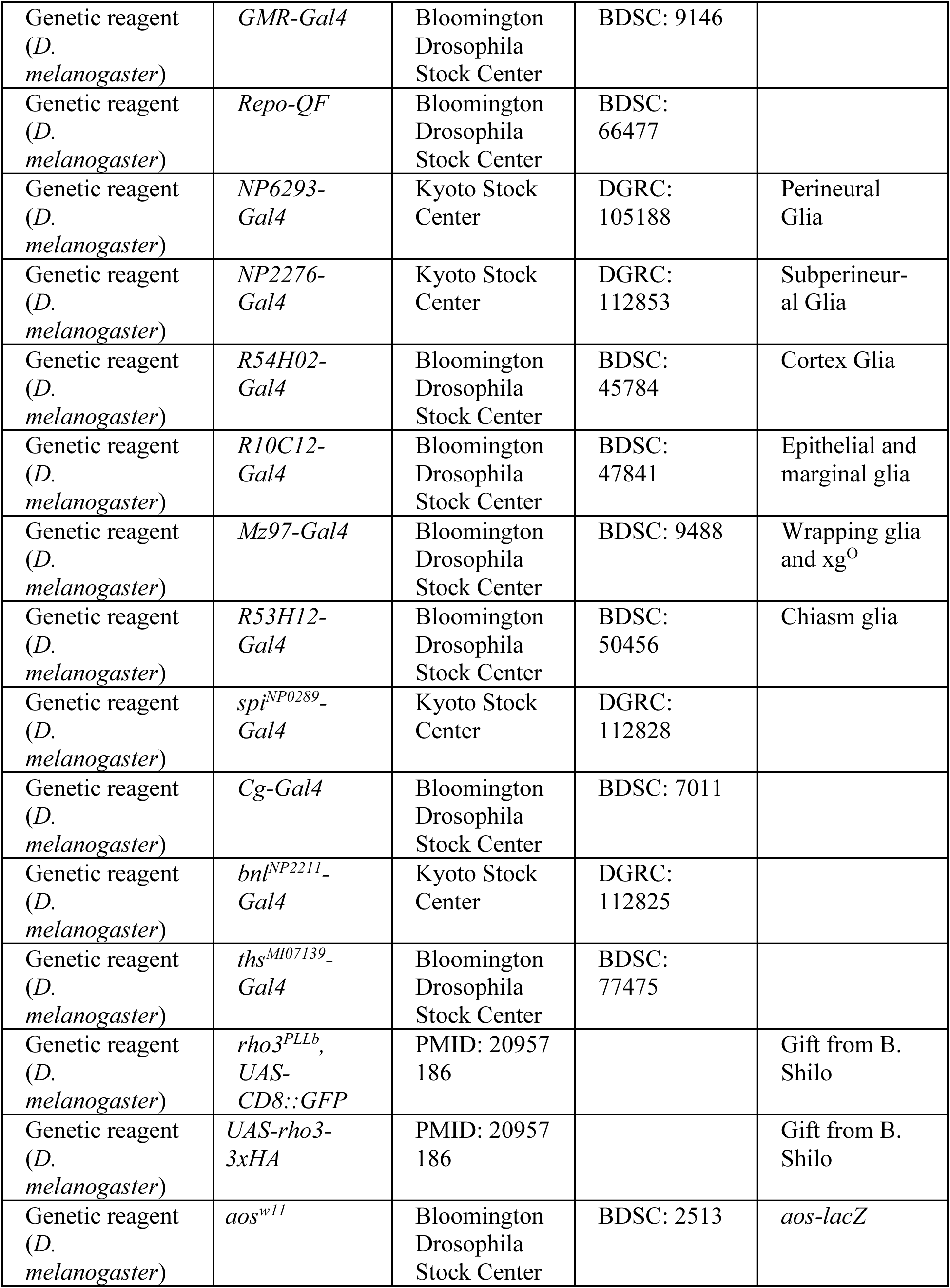
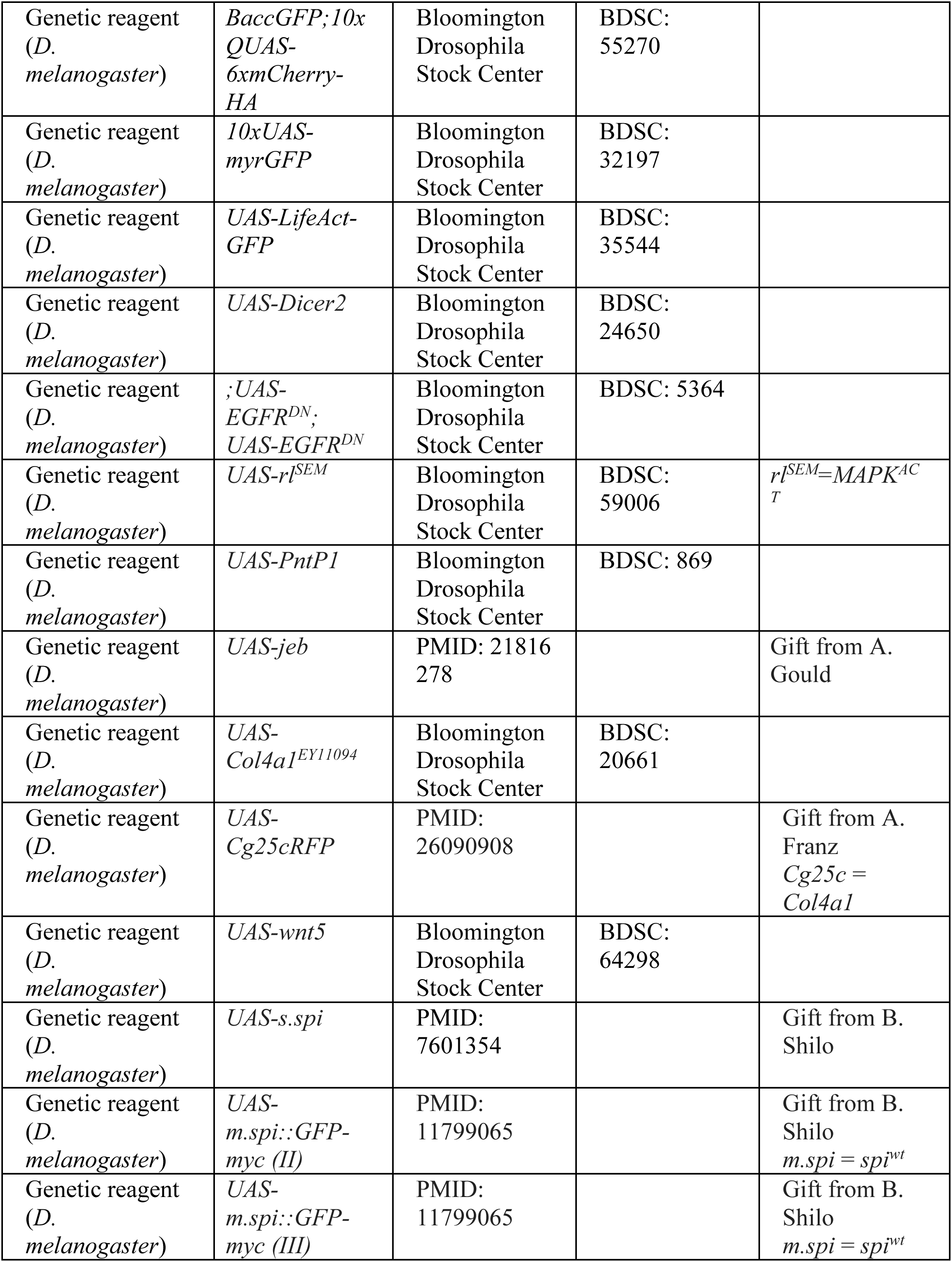
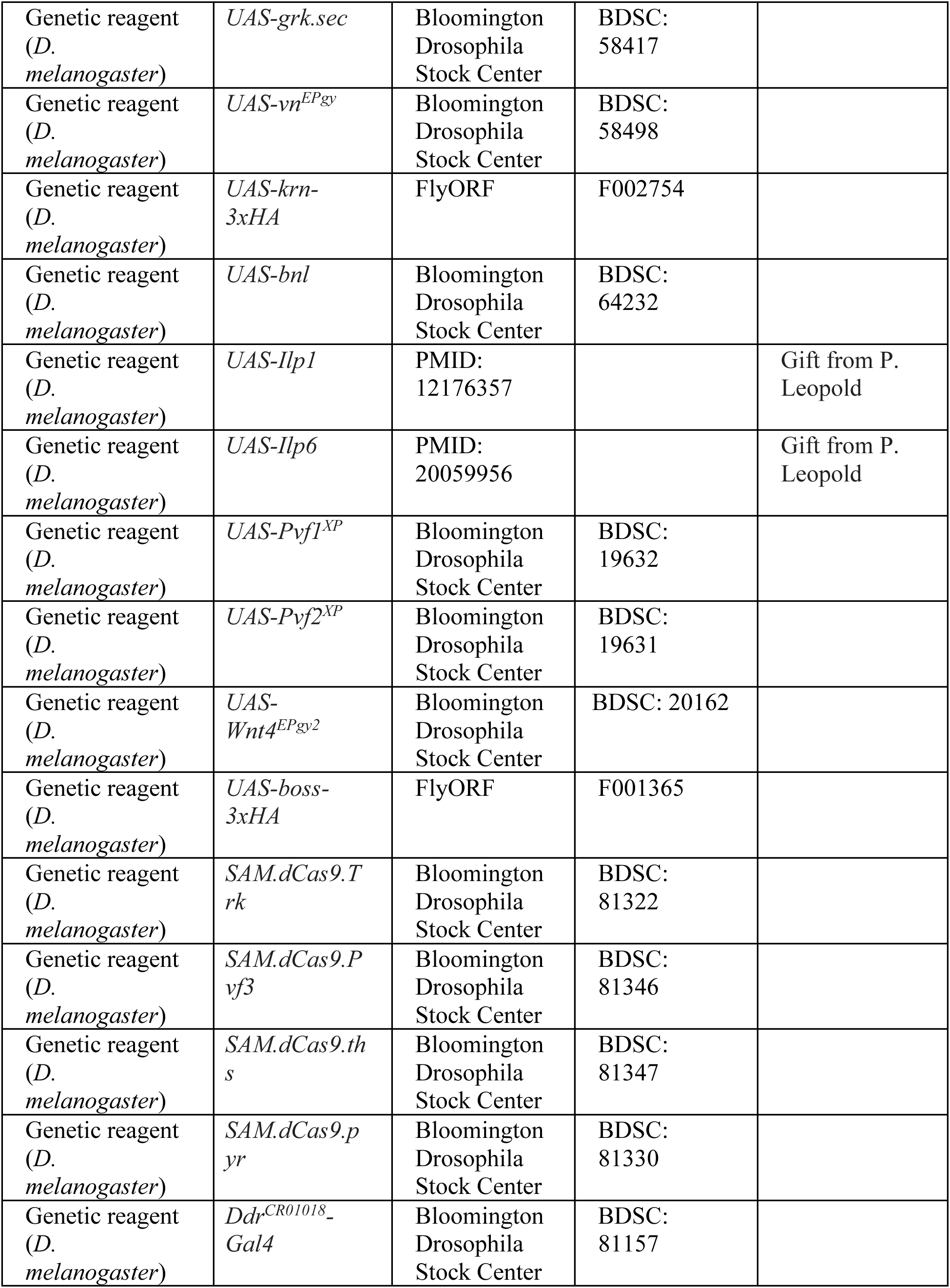
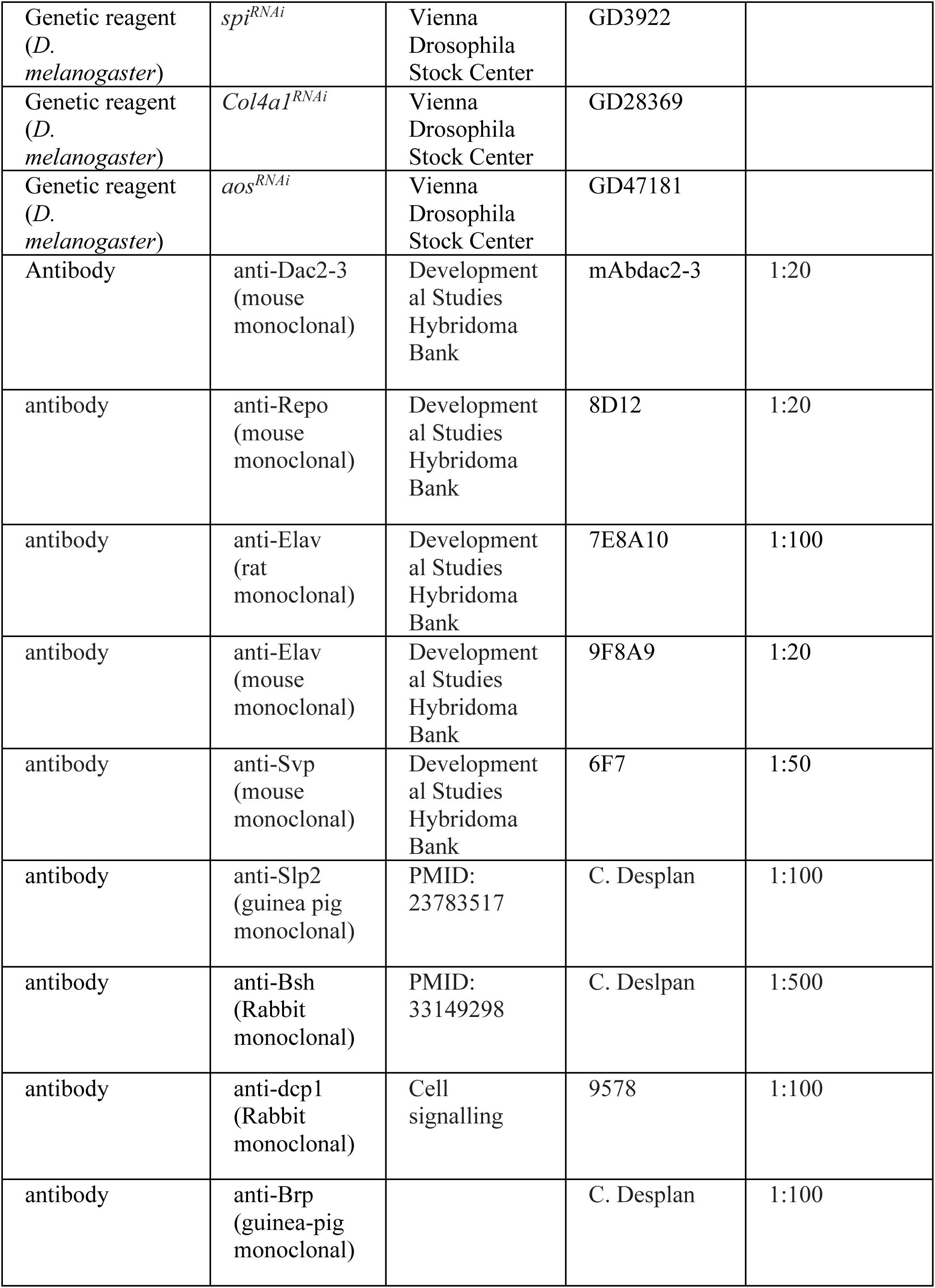
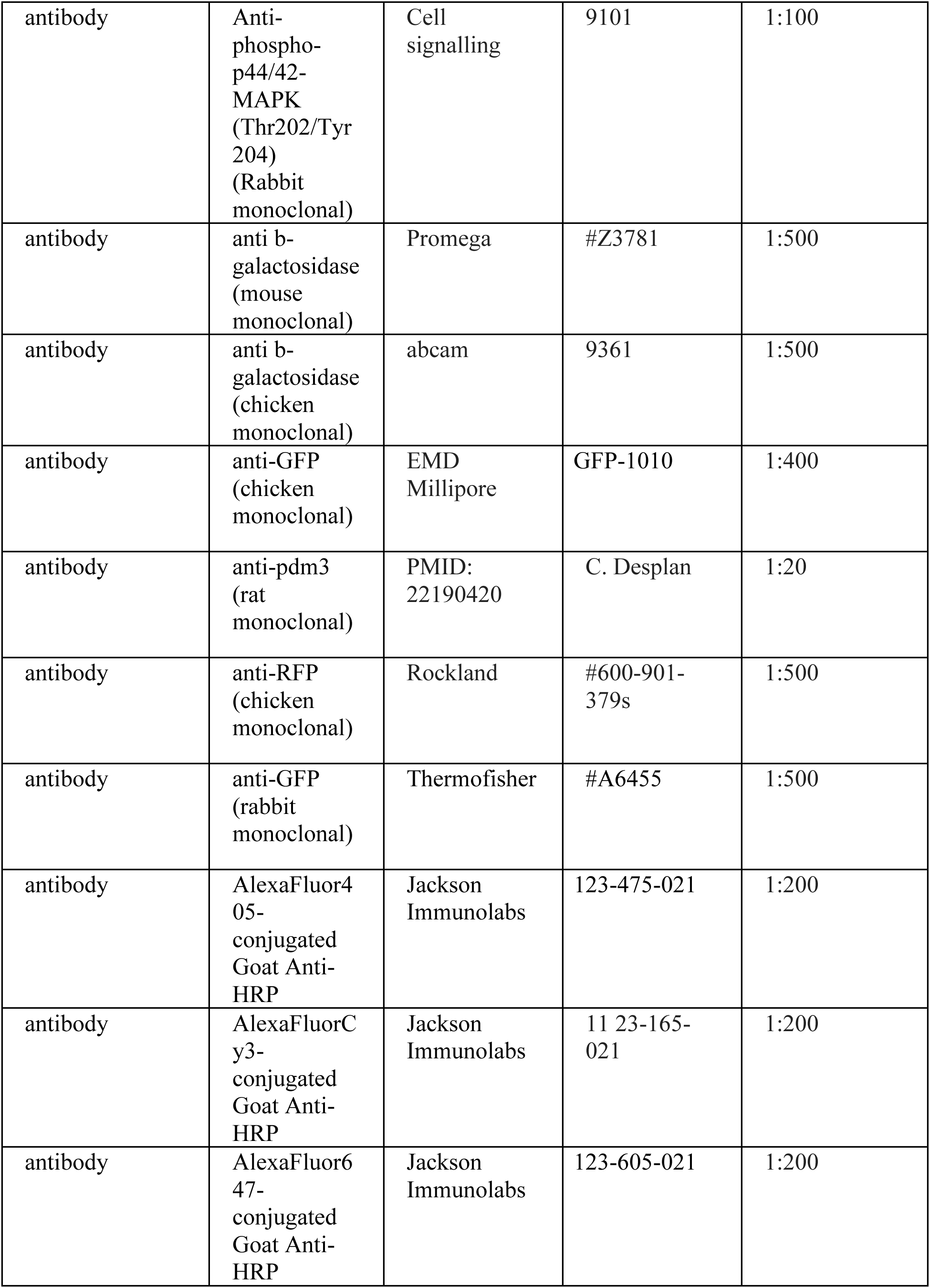
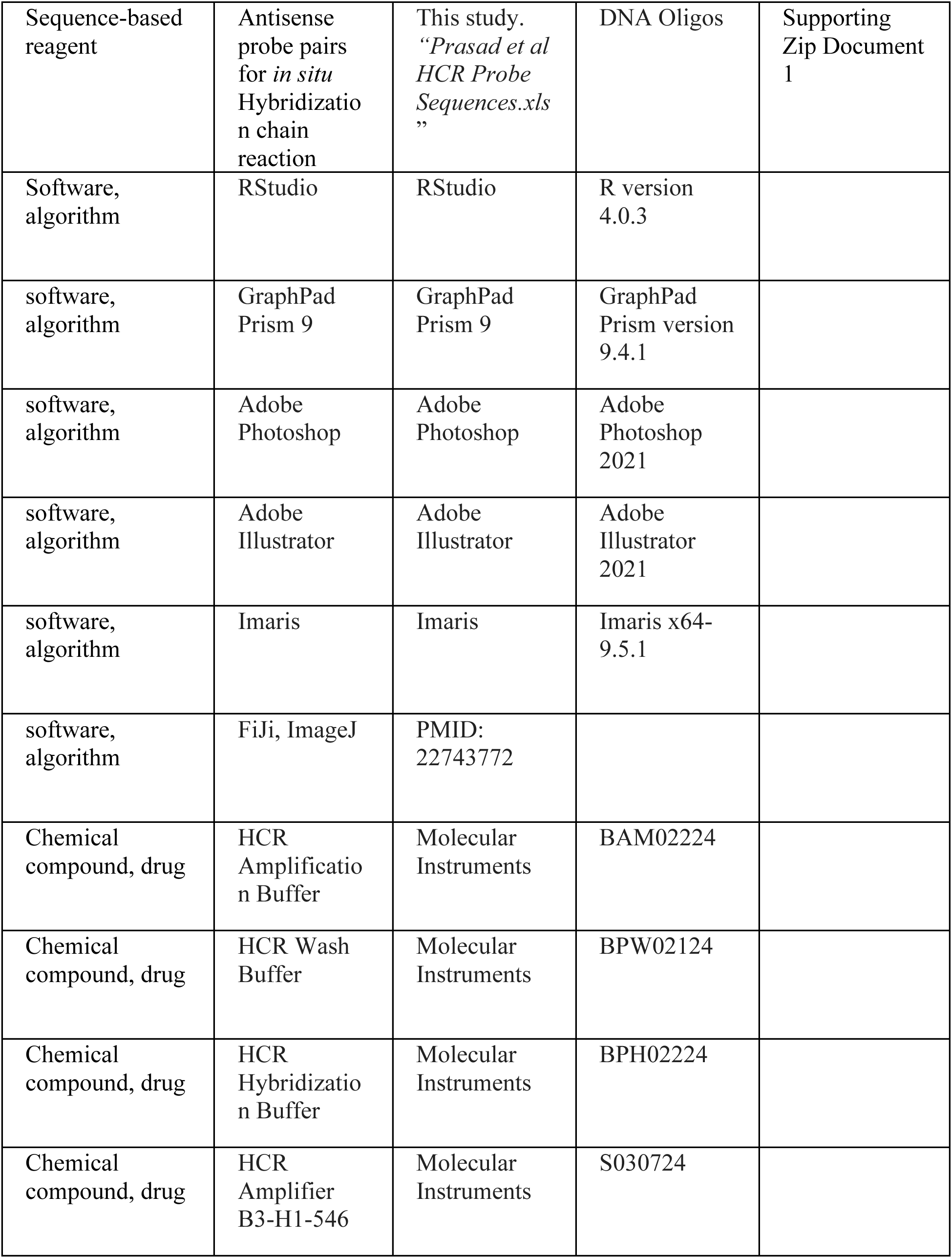
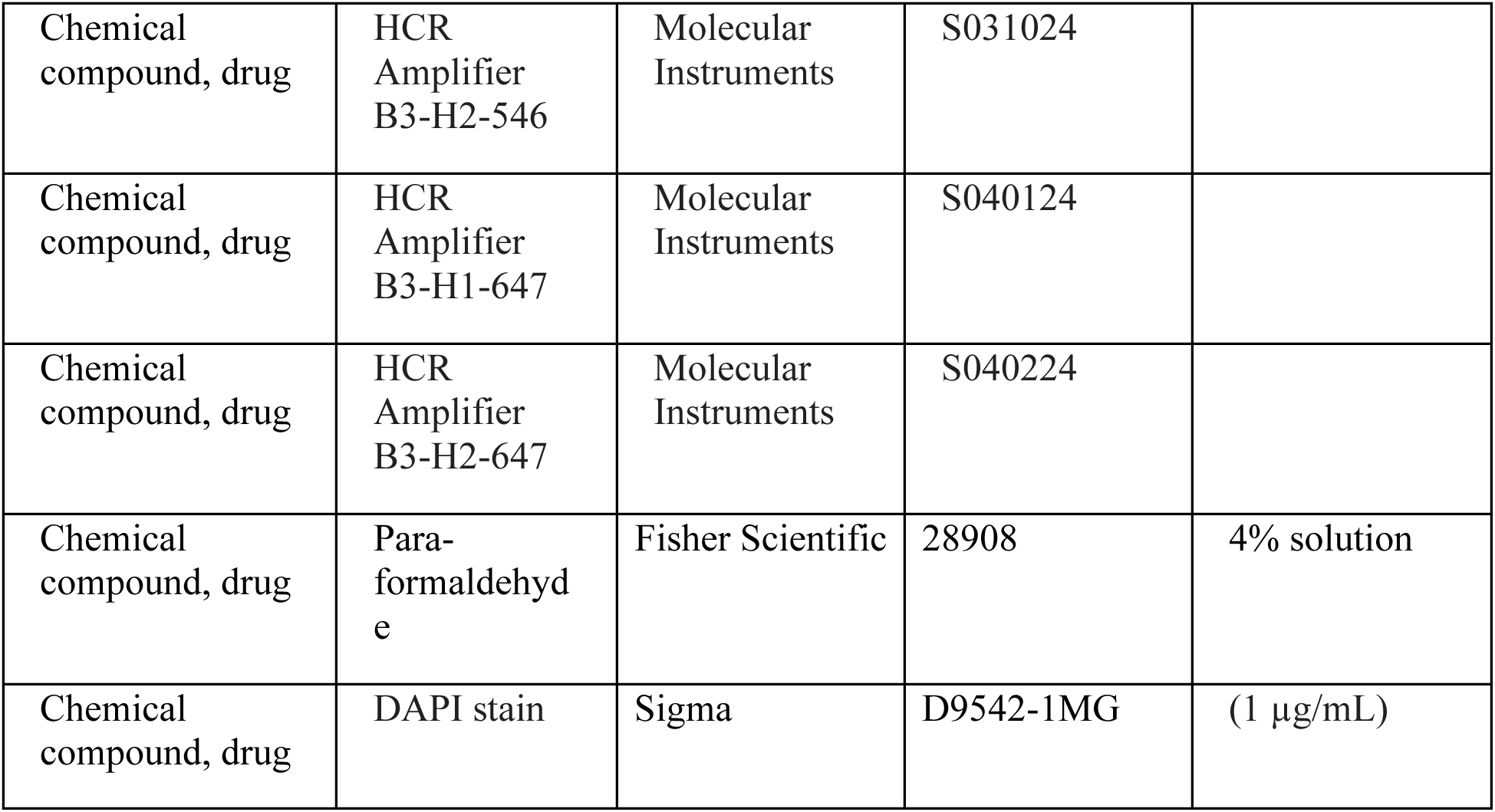

### Drosophila stocks and maintenance

*Drosophila melanogaster* strains and crosses were reared on standard cornmeal medium and raised at 25°C or 29°C or shifted from 18°C to 29°C for genotypes with temperature sensitive Gal80, as indicated in Supporting File 2.

We used the following mutant and transgenic flies in combination or recombined in this study (See Supporting File 2 for more details; {} enclose individual genotypes, separated by commas):

*{y,w,hsflp^122^; sp/Cyo; TM2/TM6B}, {y,w; sp/Cyo, Bacc-GFP; Dr/TM6C},* (from BDSC: 36349) *{ey-Gal80; sp/Cyo;}* (BDSC: 35822), *{;Gal80^ts^; TM2/TM6B}* (BDSC: 7108), *{w^1118^;; R27G05- Gal4}* (BDSC: 48073), *{w^1118^;;25A01-Gal4}* (BDSC: 49102), *{y,w; R64B07-Gal4;}* (larval L5- Gal4), {y,w; hh-Gal4/TM3} (BDSC: 67493), *{;tub-Gal80^ts^; repo-Gal4/TM6B}, {w^1118^;GMR- Gal4/Cyo;}* (BDSC: 9146), *{y,w;Pin/Cyo;repo-QF/TM6B}* (BDSC: 66477), *{y,w; NP6293- Gal4/Cyo,UAS-lacZ;}* (perineurial glia; Kyoto Stock Center: 105188), *{w; NP2276-Gal4/Cyo; }* (subperineurial glia; Kyoto Stock Center: 112853)*, {w^1118^;; R54H02-Gal4}* (cortex glia; BDSC: 45784), *{w^1118^;; R10C12-Gal4}* (epithelial and marginal glia; BDSC: 47841), *{w;Mz97-Gal4, UAS-Stinger/Cyo;}* (wrapping and xg^O^; BDSC: 9488), *{w^1118^;; R53H12-Gal4}* (chiasm glia; BDSC: 50456), *{y,w; spi^NP0289^-Gal4/Cyo, UAS-lacZ;}* (Kyoto Stock Center: 112128), *{w^1118^; Cg- Gal4;}* (BDSC: 7011), *{w;; bnl^NP2211^-Gal4}* (Kyoto Stock Center: 112825), *{w; ths^MI07139^- Gal4/Cyo; MKRS/TM6B}* (BDSC: 77475), *{;;rho3^PLLb^, UAS-CD8::GFP/TM6B}, {;UAS-rho3-3xHA;}* (gifts from B. Shilo), *{;;aos^w11^/TM6B}* (*aos-lacZ*; BDSC: 2513), *{y,w; sp/Cyo, Bacc- GFP; 10xQUAS-6xmCherry-HA}* (BDSC: 52270), *{y,w;;10xUAS-myrGFP}* (BDSC: 32197), *{;UAS-CD8::GFP;}, {;;UAS-CD8::GFP}* (gifts from C. Desplan)*, {y,w;;UAS-nls.lacZ},* (BDSC: 3956)*, {y,w; UAS-LifeAct-GFP/Cyo;}* (BDSC: 35544)*, {w^1118^;UAS-Dcr-2;}* (BDSC: 24650), *{w^1118^;;UAS-Dcr-2}* (BDSC: 24651), *{;UAS-EGFR^DN^; UAS-EGFR^DN^}* (BDSC: 5364)*, {;UAS- aop^ACT^;}* (Kyoto Stock Center: 108425), *{y,w;UAS-rl^sem^;}* (rl^sem^ = MAPK^ACT^; BDSC: 59006), *{w^1118^;;UAS-PntP1}* (BDSC: 869), *{w^1118^;UAS-aos^RNAi^;}* (VDRC47181), *{w;UAS-jeb;}* (a gift from A. Gould), *{y,w, UAS-Col4a1^EY11094^/(Cyo);}* (BDSC: 20661), *{;;UAS-Cg25c-RFP}* (Zang et al., 2015) *(Col4a1 = Cg25c)*, *{;UAS-Wnt5;}* (BDSC: 64298), *{;;UAS-s.spi}* (a gift from B. Shilo), *{;UAS-m.spi::GFP-myc;}* (a gift from B. Shilo), *{;;UAS-m.spi::GFP-myc}* (a gift from B. Shilo), *{w, UAS-grk.sec/Cyo;}* (BDSC: 58417), *{;UAS-vn^EPgy^/Cyo;}* (BDSC: 58498), *{;;UAS-krn-3xHA}* (FlyORF: F002754), *{;UAS-bnl/Cyo; MKRS/TM6C}* (BDSC: 64232), *{;UAS-Ilp1;}, {;UAS-Ilp6;}* (gifts from P. Leopold), *{w^1118^, UAS-Pvf1^XP^;;}* (BDSC: 19632), *{w^1118^; UAS- Pvf2^XP^;}* (BDSC: 19631), *{;UAS-Wnt4^EPgy2^/Cyo;}* (BDSC: 20162), *{;;UAS-boss-3xHA}* (FlyORF: F001365), *{y,sev; SAM.dCas9.Trk;}* (BDSC: 81322), *{y,sev; SAM.dCas9.Pvf3;}* (BDSC: 81346), *{y,sev; SAM.dCas9.ths;}* (BDSC: 81347), *{y,sev; SAM.dCas9.pyr;}* (BDSC: 81330), *{w^1118^; Ddr^CR01018^-Gal4;}* (BDSC: 81157).

### Immunocytochemistry, antibodies and microscopy

We dissected eye-optic lobe complexes from early pupae (0-5hrs after puparium formation; APF) in 1X phosphate-buffered saline (PBS), fixed in 4% formaldehyde for 20 minutes, blocked in 5% normal donkey serum and incubated in primary antibodies diluted in block for 2 nights at 4°C. Samples were then washed in 1X PBS with 0.5% TritonX (PBSTx), incubated in secondary antibodies diluted in block, washed in PBSTx and mounted in SlowFade (Life Technologies). When performing phospho-MAPK stains, dissections were performed in a phosphatase inhibitor buffer as detailed in (Amoyel et al., 2016).

We used the following primary antibodies in this study: mouse anti-Dac^2-3^ (1:20, Developmental Studies Hybridoma Bank; DSHB), mouse anti-Repo (1:20, DSHB), rat anti-Elav (1:100, DSHB), mouse anti-Elav (1:20, DSHB), rabbit anti-Dcp-1 (1:100; Cell Signalling #9578), chicken anti- GFP (1:400; EMD Millipore), mouse anti-Svp (1:50, DSHB), rabbit anti-Slp2 (1:100; a gift from C. Desplan), rabbit-Bsh (1:500; a gift from C. Desplan), Rat anti-Pdm3 (1:1000; a gift from C. Desplan), guinea pig anti-Brp (1:100; a gift from C. Desplan), rabbit anti-Phospho-p44/42 MAPK (Erk1/2) (Thr202/Tyr204) (1:100, Cell Signaling #9101), chicken anti-RFP (1:500; Rockland #600-901-379s), mouse anti b-galactosidase (1:500; Promega # Z3781), chicken anti b-galactosidase (1:500; abcam #9361), rabbit-anti-GFP (1:500; Thermofisher #A6455), AlexaFluor405 conjugated Goat Anti-HRP (1:100; Jackson Immunolabs), AlexaFluor405-, Cy3-, or AlexaFluor647- conjugated Goat Anti-HRP (1:200; Jackson Immunolabs). Secondary antibodies were obtained from Jackson Immunolabs or Invitrogen and used at 1:800. Images were acquired using Zeiss 800 and 880 confocal microscopes with 40X objectives.

### In-situ Hybridisation Chain reaction

<u>To determine if *spi, Col4a1 and Ddr* transcripts were present in the xg^O^, we performed HCR as</u> <u>detailed in</u> Duckhorn et al. (2022)<u>. We designed 20-21 probe pairs against target genes, excluding regions of strong similarity to other transcripts, with corresponding initiator sequences</u> <u>for amplifiers B3</u> (Choi et al., 2018)<u>. HCR probes (sequences included as source data</u> See Supporting Zip Document 1<u>) were purchased as DNA Oligos from ThermoFisher (100µm in</u> <u>water and frozen).</u>

Eye-optic lobe complexes were dissected, fixed and washed as detailed above. Samples were incubated in Probe Hybridisation Buffer for 30 minutes at 37°C followed by incubation with probes (0.01µM) at 37°C overnight. The samples were then washed four times for 15 minutes each with probe wash buffer at 37°C followed by two washes for 5 minutes each with 5X saline sodium citrate solution (20XSSCT solution in distilled water-- 58.44g/mol sodium chloride, 294.10g/mol 560 sodium citrate, pH adjusted to 7 with 14N hydrochloric acid, with 0.001% Tween 20) at room temperature. Samples were then incubated with amplification buffer for 10 minutes at room temperature. 12 pmol of hairpins H1 and H2 were snap-cooled (95°C for 90 seconds and then cooled to room-temperature for 20 mins) separately to avoid oligomerisation. The snap-cooled hairpins were then added to the samples in the amplification buffer (protected from light) and incubated overnight at room temperature. The samples were then washed with 5XSSCT for 15 minutes before being incubated in darkness with 1:15 dilution of DAPI (Sigma D9542) for 90 minutes. Samples were washed with 1xPBS for 30 minutes and then mounted as detailed above.

### Quantification and Statistical analyses

We used Fiji-ImageJ (Schindelin et al., 2012) or Imaris (version x64-9.5.1) to process and quantify confocal images as described below. We used Adobe Photoshop and Adobe Illustrator software to prepare figures. We used GraphPad Prism8 to perform statistical tests. In all graphs, whiskers indicate the standard error of the mean (SEM).

#### Dcp-1 quantifications

We used the surfaces tool in Imaris to manually select the lamina region (based on Dac expression). We then used the spots tool to identify Dcp-1 positive cells (cell diameter = 5µm) within the selected region using the default thresholding settings, and plotted these values normalised to the volume of the selected lamina region in GraphPad Prism8.

#### Cell-type quantifications

*LPCs per column:* Column number was identified by counting HRP-labelled photoreceptor axon bundles. We considered the youngest column located adjacent to the lamina furrow to be the first column, with column number (age) increasing towards the posterior (right) of the furrow. We counted the number of Dac+ cells per column by quantifying 10 optical slices (step size = 1µm) located centrally in the lamina.

#### Control vs. Lamina^ts^>PntP1

We quantified the lamina neuron types per column using the following markers to identify L-neuron types: Elav+ and Slp2+ cells were counted as L1-L3s; Elav+ and Bsh+ cells were counted as L4s and Elav+, Bsh+ and Slp2+ cells were counted as L5s. We quantified 10 optical slices (step size = 1µm) located centrally in the lamina. Column number was identified by counting HRP-labelled photoreceptor axon bundles. These quantifications were done blind.

#### Ligand receptor screen

We quantified the number of L5s based on Elav expression in the proximal lamina. Column number was identified by counting HRP-labelled photoreceptor axon bundles. We quantified 30 optical slices (step size = 1µm) located centrally in the lamina.

#### Ligand over-expression quantifications

We quantified the number of L-neuron types per column using Elav, Bsh and Slp2. We quantified 30 optical slices (step size = 1µm) located centrally in the lamina. Column number was identified by counting HRP-labelled photoreceptor axon bundles.

#### Spi and Col4a1 Probe Intensity Quantifications

In Fiji-ImageJ we used the free hand selection tool to draw a region of interest (ROI) around the xg^O^ (marked by the *xg^O^>CD8::GFP*). We then measured the mean fluorescence intensity of *spi* and *Col4a1* transcripts labelled by HCR within each ROI. We quantified 30 optical slices (step size = 1µm) located centrally in the lamina and then plotted the average for each optic lobe.

#### Number of xg^O^

We quantified the number of xg^O^ (Figure S1Q) by manually counting the number of Repo positive nuclei within LifeAct-GFP positive xg^O^ per 40µm optical section in Fiji-ImageJ. We used a step size of 1mm while acquiring the z-stacks and centred each 40µm optical section in the middle of the lamina using photoreceptor axons (HRP), and the lobula plug (Dac expression) as landmarks. Quantifications were performed blind.

#### Length of xg^O^ processes

We quantified the lengths of the fine glial processes that extend distally from the xg^O^ towards the lamina plexus (Figure S1O,P,R) by using the straight-line selection and measuring tools in Fiji-ImageJ to measure xg^O^ process lengths in a 10mm optical section centred in the middle of the lamina. Quantifications were performed blind.

#### Quantifications of nuclear to cytoplasmic dpMAPK mean fluorescence intensity

Using Fiji we manually drew regions of interest (ROIs) with the free hand selection tool around the xg^O^ nucleus (based on Repo) and LPCs in the most proximal row of the lamina (based on Dac expression) and added these to the ROI manager. We then enlarged the ROIs (Edit>Selection>Enlarge) by 3.00 pixel units to include the cytoplasm. We then used the XOR function in the ROI Manager to only select the cytoplasm of the xg^O^. We then measured the MFI of dpMAPK in the nucleus and the cytoplasm of the xg^O^ in 20 centrally located optical slices (corresponding to 20µm) for each optic lobe. We plotted the nuclear:cytoplasmic ratios of dpMAPK MFI in GraphPad Prism8.

#### aos-lacZ intensity quantifications

Using Fiji we manually drew regions of interest around L5s (based on Slp2 + Bsh co-expression) in each column. We then measured the mean fluorescence intensity of b-Gal in the ROIs. We quantified 10 optical slices (step size = 1µm) for each optic lobe and plotted the average values as a function of column (age).

**Figure 1-figure supplement 1:**
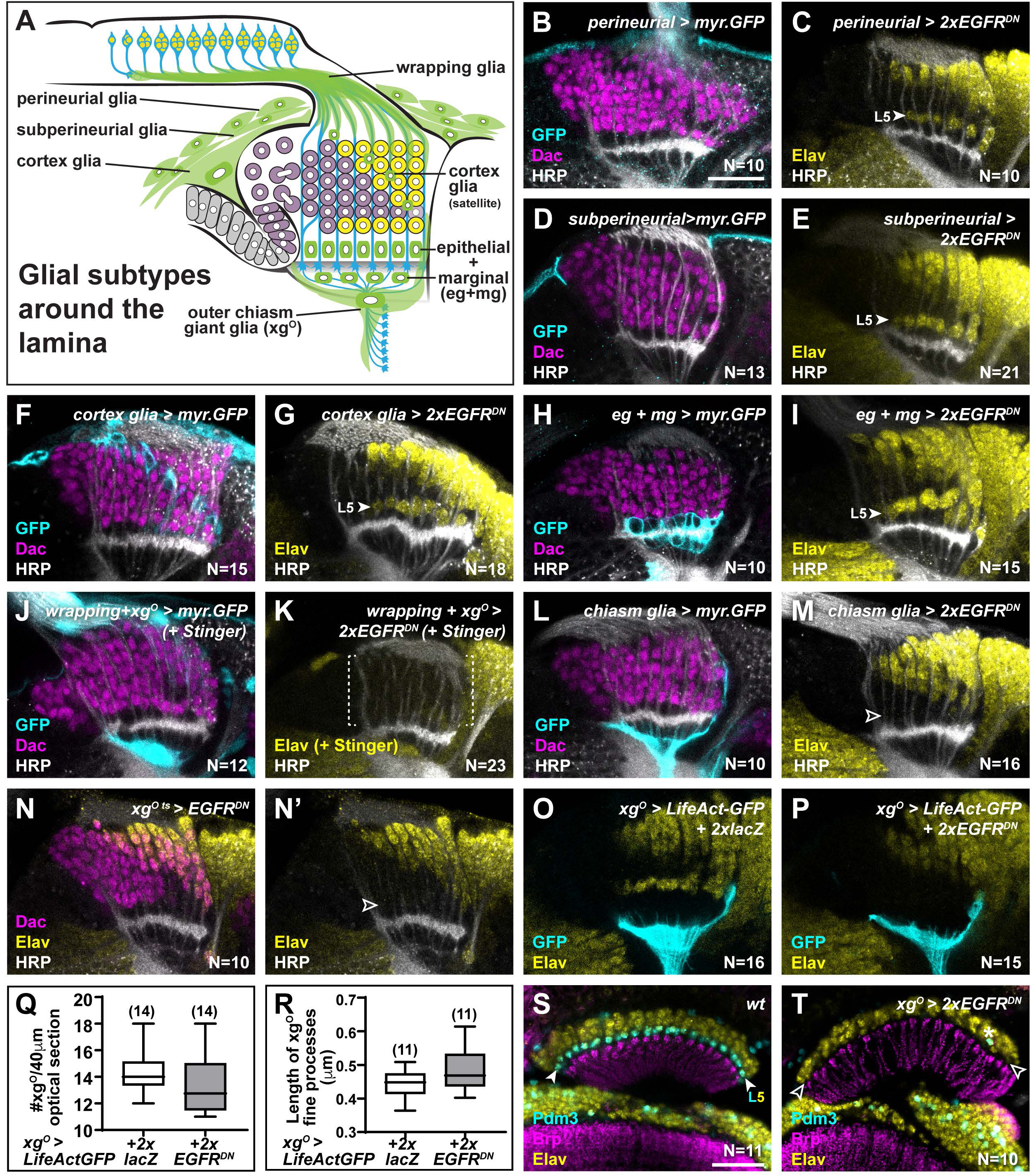
A Gal4 screen identifies xg^O^ as the glial sub-type that regulates L5 neuronal differentiation. **(A)** Schematic of the developing lamina and associated glial types (green; labelled). **(B)** A perineurial glia specific Gal4 drives expression of myr.GFP stained for GFP (cyan), Dac (magenta) and HRP (white). **(C)** Perineurial glia specific expression of EGFR^DN^ stained for Elav (yellow) and HRP (white). L5 differentiation was not affected. **(D)** A subperineurial glia specific Gal4 drives expression of myr.GFP stained for GFP (cyan), Dac (magenta) and HRP (white). **(E)** Suberineurial glia specific expression of EGFR^DN^ stained for Elav (yellow) and HRP (white). L5 differentiation was not affected. **(F)** A cortex glia specific Gal4 drives expression of myr.GFP stained for GFP (cyan), Dac (magenta) and HRP (white). **(G)** Cortex glia specific expression of EGFR^DN^ stained for Elav (yellow) and HRP (white). L5 differentiation was not affected. **(H)** An epithelial and marginal glia (eg+mg) specific Gal4 drives expression of myr.GFP stained for GFP (cyan), Dac (magenta) and HRP (white). **(I)** Epithelial and marginal glia specific expression of EGFR^DN^ stained for Elav (yellow) and HRP (white). L5 differentiation was not affected. **(J)** A wrapping glia and xg^O^ specific Gal4 drives expression of myr.GFP stained for GFP (cyan), Dac (magenta) and HRP (white). **(K)** Wrapping glia and xg^O^ specific expression of EGFR^DN^ stained for Elav (yellow) and HRP (white). L1-L4 and L5 differentiation were disrupted as observed by the lack of Elav+ cells in the lamina. **(L)** A chiasm glia (xg^O and^ xg^inner)^ specific Gal4 drives expression of myr.GFP stained for GFP (cyan), Dac (magenta) and HRP (white). **(M)** Chiasm glia specific expression of EGFR^DN^ stained for Elav (yellow) and HRP (white). L1- L4 differentiation proceeded normally but L5 differentiation was disrupted as observed by the lack of Elav+ cells in the proximal lamina. **(N)** Gal80^ts^-restricted Gal4 expression in xg^O^, driving EGFR^DN^ during lamina development (See Supporting File 1) stained for Dac (magenta), Elav (yellow) and HRP (white). L5 neurons were dramatically reduced. **(O,P)** LifeAct-GFP expression driven in xg^O^ in **(O)** controls and **(P)** when 2 copies of EGFR^DN^ are coexpressed. In both conditions, the fine processes from the xg^O^ are present. **(Q)** Quantification of xg^O^ numbers in control *xg^O^>LifeAct-GFP+2xlacZ* and *xg^O^>LifeAct-GFP + 2xEGFR^DN^.* P>0.05; Mann-Whitney U test. Ns indicated in parentheses. **(R)** Quantification of the length of xg^O^ fine processes in control *xg^O^>LifeAct-GFP+2xlacZ* and *xg^O^>LifeAct-GFP + 2xEGFR^DN^.* P>0.05; Unpaired t test. Ns indicated in parentheses. **(S)** Wildtype adult optic lobe stained for Pdm3 (L5 marker) (Tan et al., 2015), Bruchpilot (Brp; marks neuropils) and Elav (yellow). **(T)** *xg^O^>2xEGFR^DN^* adult optic lobe stained for Pdm3 (L5 marker) (Tan et al., 2015), Bruchpilot (Brp; marks neuropils) and Elav (yellow). Pdm3+ cells (L5s) are reduced dramatically. Scale bar = 20µm. For all quantifications boxes indicate the lower and upper quartiles; the whiskers represent the minimum and maximum values; the line inside the box indicates the median).

**Figure 3-figure supplement 1:**
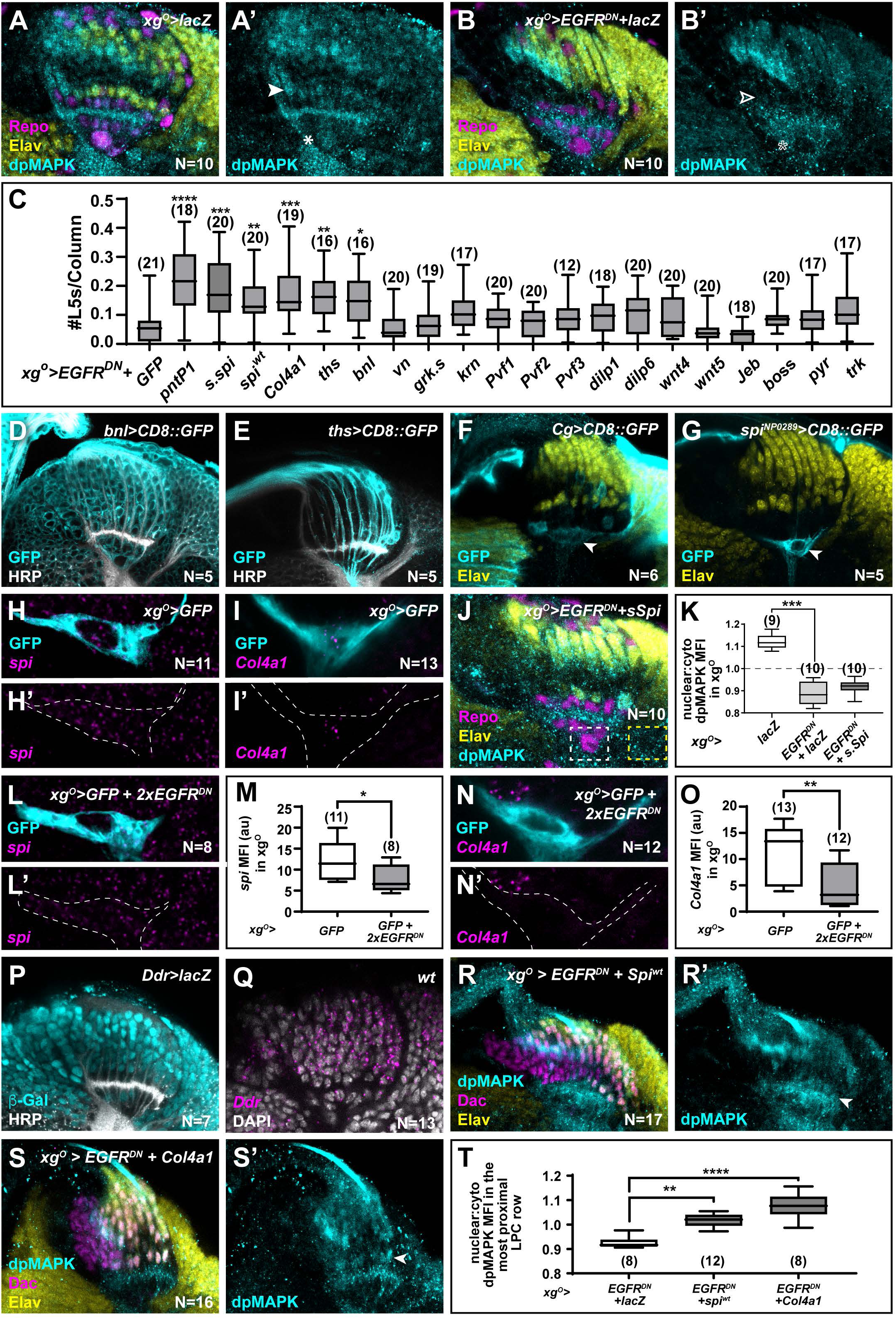
Multiple xg^O^ secreted ligands activate MAPK signalling to drive L5 neuronal differentiation. **(A,B)** Optic lobes stained for Elav (yellow), Repo (magenta) and dpMAPK (cyan) in **(A)** *xg^O^>lacZ* controls and **(B)** with EGFR^DN^ and lacZ expressed in xg^O^. dpMAPK levels decreased in the xg^O^ (indicated by asterisk) and in cells in the proximal row of the lamina (indicated by arrowhead) when compared with *xg^O^>lacZ* controls. **(C)** Quantification of the number of L5s/column (based on Elav expression) when different ligands that can activate MAPK signalling were co-expressed with EGFR^DN^ in the xg^O^. (P*<0.05; P**<0.01; P***<0.0005; P****<0.0001; one-way ANOVA with Dunn’s multiple comparison test. Ns indicated in parentheses). **(D)** *bnl>CD8::GFP* showed GFP (cyan) expression in all cells in the optic lobe; HRP (white). **(E)** *ths>CD8::GFP* showed GFP (cyan) expression in photoreceptors; HRP (white). **(F)** *Collagen>CD8::GFP* drove GFP (cyan) expression in xg^O^ (arrowhead); Elav (yellow). **(G)** *spi^NP0289^>CD8::GFP* drove GFP (cyan) expression in xg^O^ (arrowhead); Elav (yellow). **(H, I)** *xg^O^>GFP* lobes stained for GFP (cyan) and **(H)** *spi* mRNA (magenta) and **(I)** *Col4a1* mRNA (magenta) by *in situ* hybridisation chain reaction (HCR). **(J)** *xg^O^>EGFR^DN^+s.spi* lobes stained for Elav (yellow), Repo (magenta) and dpMAPK (cyan). Inset shows a magnified view of the xg^O^ nucleus. **(K)** Quantifications of nuclear:cytoplasmic ratios of dpMAPK MFI in the xg^O^ in indicated genotypes (P<0.0005, one-way ANOVA with Dunn’s multiple comparisons test. Ns indicated in parentheses) **(L)** *spi* mRNA (magenta) detected by HCR in *xg^O^>GFP+2xEGFR^DN^* lobes. **(M)** Quantification of *spi* MFI (arbitrary units) for (H and L). (P<0.05; Mann-Whitney U test.) **(N)** *Col4a1* mRNA (magenta) detected by HCR in *xg^O^>GFP+2xEGFR^DN^* lobes. **(O)** Quantification of *Col4a1* MFI (arbitrary units) for (I and N) (P<0.005; Mann-Whitney U test). **(P)** *Ddr>lacZ* showed b-Galactosidase (b-Gal; cyan) expression in the lamina; HRP (white). **(Q)** *Ddr* mRNA (magenta) detected by HCR in w*ild-type* lobes; DAPI (white). **(R,S)** Lobes stained for Dac (magenta), Elav (yellow) and dpMAPK (cyan) when **(R)** Spi^wt^ is coexpressed with EGFR^DN^ in xg^O^ or **(S)** Col4a1 is coexpressed with EGFR^DN^ in xg^O^. Arrowheads indicate Elav+ cells in the most proximal row. **(T)** Quantifications of nuclear:cytoplasmic ratios of dpMAPK MFI in the most proximal row of LPCs in indicated genotypes (P**<0.005. P****<0.0001; one-way ANOVA with Dunn’s multiple comparisons test). Scale bar = 20µm. For all quantifications boxes indicate the lower and upper quartiles; the whiskers represent the minimum and maximum values; the line inside the box indicates the median).

**Figure 3-figure supplement 2:**
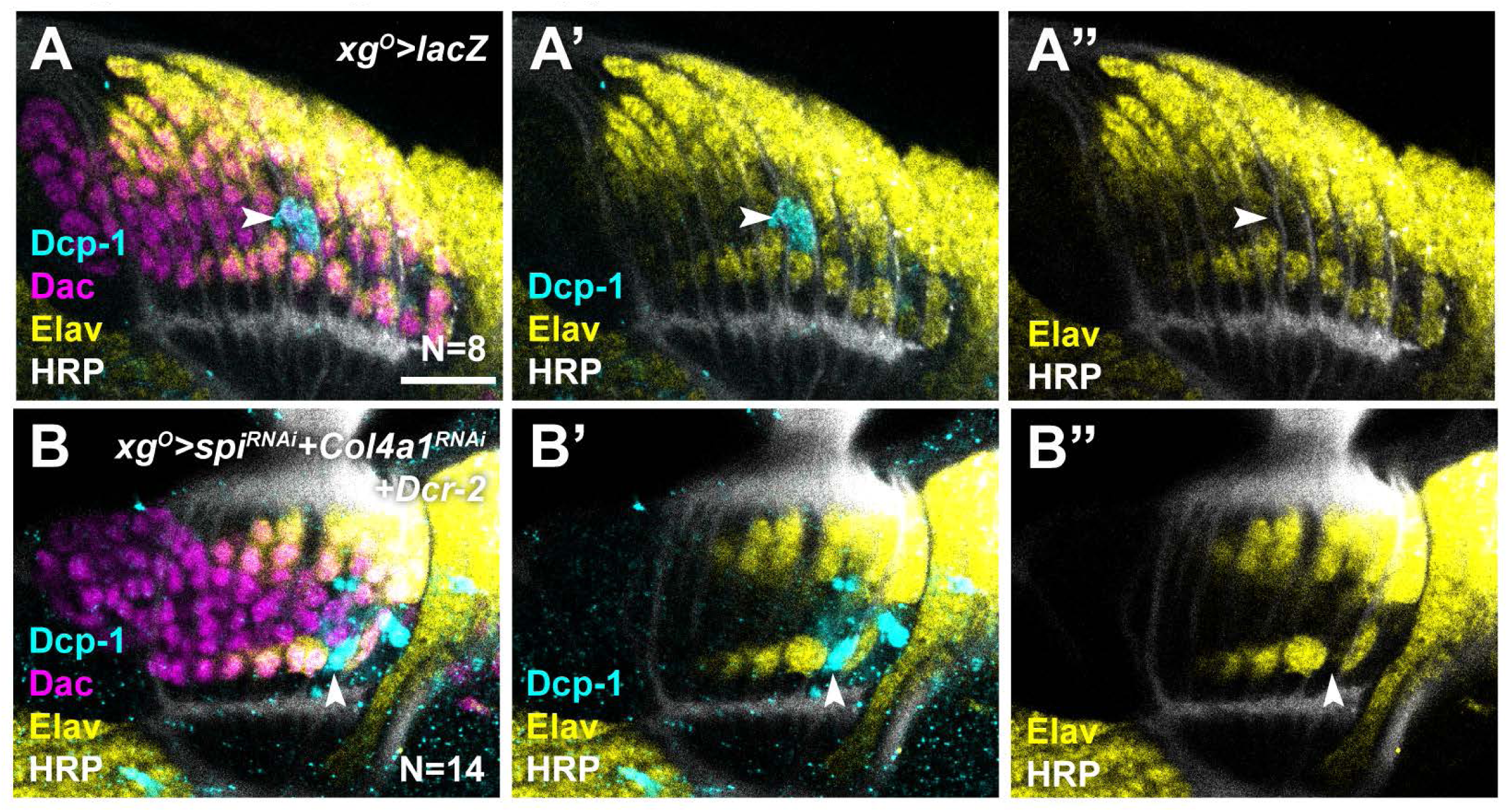
Spi and Col4a1 from xg^O^ promote cell survival in proximal LPCs. **(A)** *xg^O^>lacZ* lobes stained for Dcp-1 (cyan), Dac (magenta), Elav (yellow) and HRP (white). Dcp-1 positive cells (indicated by arrowhead) were located between L4s and L5s and corresponds to ‘extra’ LPCs which undergo apoptosis. **(B)** *xg^O^>Spi^RNAi^+Col4a^RNAi^+ Dcr-2* lobes stained for Dcp-1 (cyan), Dac (magenta), Elav (yellow) and HRP (white). Dcp-1 positive cells were observed in the proximal row of L5s (indicated by arrowhead) which were never observed in controls.

**Figure 4-figure Supplement 1:**
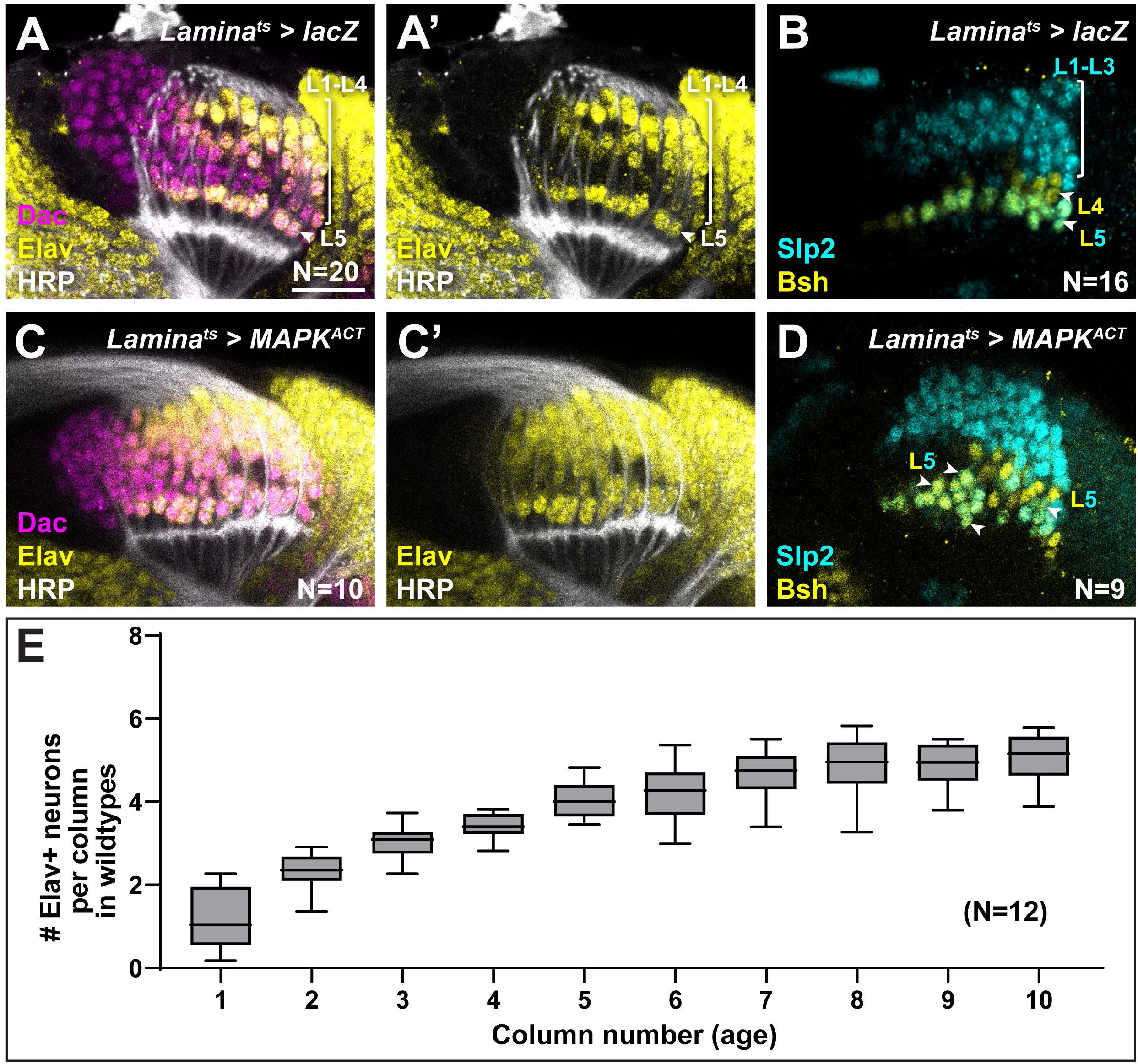
Hyperactivating MAPK in the lamina drives ectopic L5 differentiation. **(A,B)** Control *Lamina^ts^>lacZ* optic lobes stained for **(A)** Dac (magenta), HRP (white) and Elav (yellow), and **(B)** and L-neuron type specific markers Slp2 (cyan) and Bsh (yellow). **(C,D)** *Lamina^ts^>MAPK^ACT^* optic lobes stained for **(C)** Dac (magenta), HRP (white) and Elav (yellow), and **(D)** L-neuron type specific markers Slp2 (cyan) and Bsh (yellow). Ectopic Slp2 and Bsh co-expressing cells (L5s) were observed (arrowheads). **(E)** Quantification of the number of Elav+ cells per lamina column as a function of column number (age) in wildtype animals. Columns were fully differentiated (5 Elav+ cells) by column 7. Boxes indicate the lower and upper quartiles; the whiskers represent the minimum and maximum values; the line inside the box indicates the median). Scale bar = 20µm.

**Figure 5-figure supplement 1:**
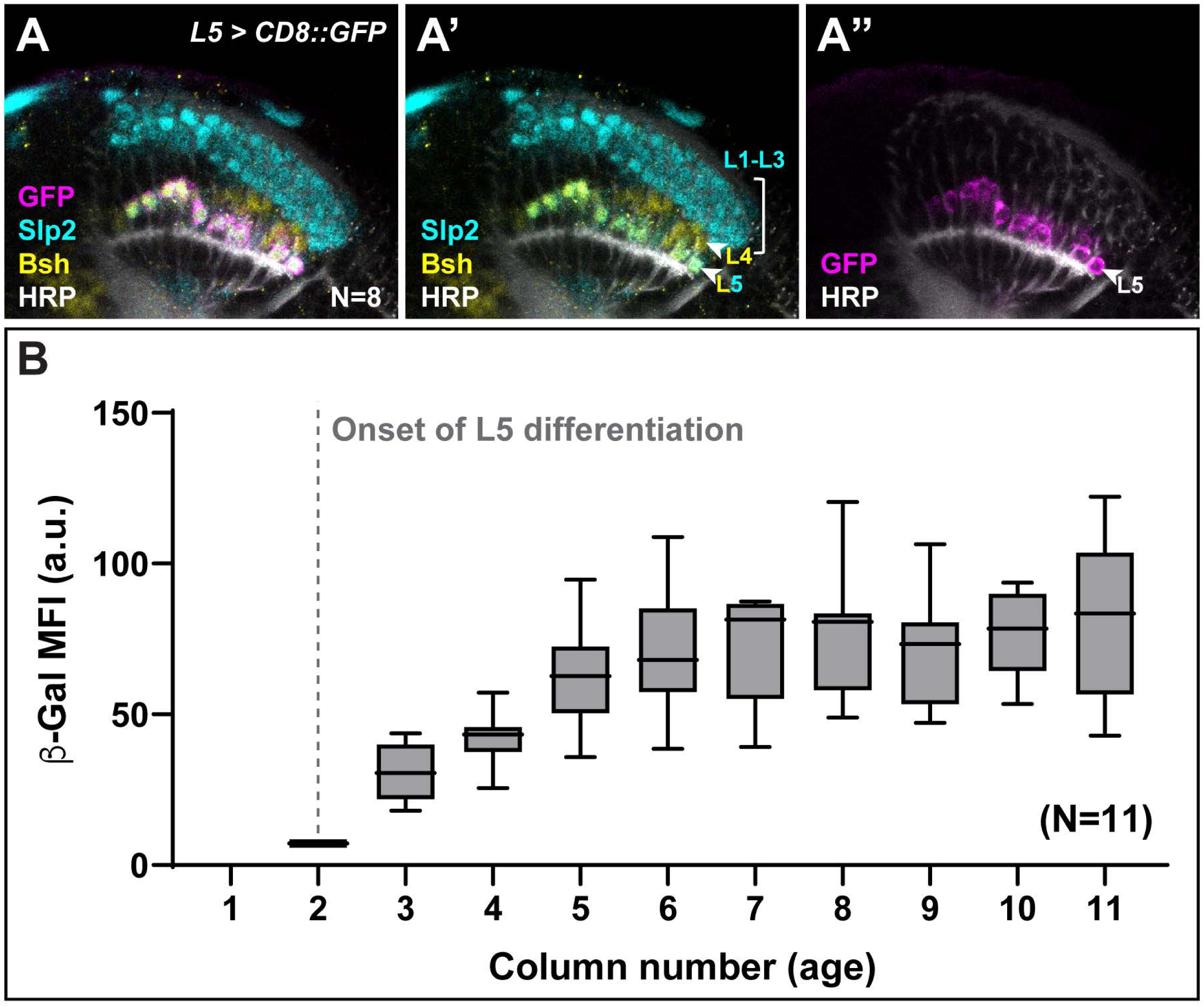
Aos expression is delayed in younger L5s. **(A)** An L5 specific driver was used to drive the expression of GFP (magenta) in the lamina; HRP (white) and L-neuron type specific markers Slp2 (cyan) and Bsh (yellow). **(B)** b-Gal MFI quantifications in the proximal row of L5s as a function of column number (age) in *aos-lacZ* lobes. b-Gal MFI is low in young columns and increases in older columns (from column 5). Boxes indicate the lower and upper quartiles; the whiskers represent the minimum and maximum values; the line inside the box indicates the median). Scale bar = 20µm.

**Supplementary File 1: Table summarising the results from the glial-Gal4 screen (Figure 1** **B,C,** **Figure 1- figure supplement 1B-N) to identify the glial type that regulates L5 development.**

**Supplementary File 2: Table listing all genotypes and experimental conditions used by figure panel.** (Note that only female genotypes are listed though both sexes were included in our analyses)

**Supporting zip document 1: Excel file containing all the probe sequences used for *in situ* hybridisation chain reaction in this study**

## References

Akagawa H, Hara Y, Togane Y, Iwabuchi K, Hiraoka T. 2015. The role of the effector caspases drICE and dcp-1 for cell death and corpse clearance in the developing optic lobe in Drosophila. Dev Biol 1–15. doi:10.1016/j.ydbio.2015.05.013

Amoyel M, Anderson J, Suisse A, Glasner J, Bach EA. 2016. Socs36E Controls Niche Competition by Repressing MAPK Signaling in the Drosophila Testis. PLoS Genet 12:e1005815. doi:10.1371/journal.pgen.1005815

Anllo L, DiNardo S. 2022. Visceral mesoderm signaling regulates assembly position and function of the Drosophila testis niche. Dev Cell 57:1009–1023.e5. doi:https://doi.org/10.1016/j.devcel.2022.03.009

Apitz H, Salecker I. 2014. A challenge of numbers and diversity: neurogenesis in the Drosophila optic lobe. J Neurogenet 28:1–35. doi:10.3109/01677063.2014.922558

Ballif BA, Blenis J. 2001. Molecular Mechanisms Mediating Mammalian Mitogen-activated Protein Kinase (MAPK) Kinase (MEK)-MAPK Cell Survival Signals. Cell Growth Differ 12:397–408.

Bergmann A, Agapite J, McCall K, Steller H. 1998. The Drosophila gene hid is a direct molecular target of Ras-dependent survival signaling. Cell 95:331–41.

Bergmann A, Tugentman M, Shilo BZ, Steller H. 2002. Regulation of cell number by MAPK- dependent control of apoptosis: A mechanism for trophic survival signaling. Dev Cell 2:159–170. doi:10.1016/S1534-5807(02)00116-8

Bostock MP, Prasad A, Donoghue A, Fernandes VM. 2022. Photoreceptors generate neuronal diversity in their target field through a Hedgehog morphogen gradient in Drosophila. Elife 11:e78093. doi:10.1101/2022.02.21.481306

Chen F, Krasnow MA. 2014. Progenitor Outgrowth from the Niche in Drosophila Trachea Is Guided by FGF from Decaying Branches. Science (80-) 343:186–189. doi:10.1126/science.1241442

Chen J, Xu N, Huang H, Cai T, Xi R. 2016. A feedback amplification loop between stem cells and their progeny promotes tissue regeneration and tumorigenesis. Elife 5:e14330. doi:10.7554/eLife.14330

Choi HMT, Calvert CR, Husain N, Huss D, Barsi JC, Deverman BE, Hunter RC, Kato M, Lee SM, Abelin ACT, Rosenthal AZ, Akbari OS, Li Y, Hay BA, Sternberg PW, Patterson PH, Davidson EH, Mazmanian SK, Prober DA, Rijn M Van De, Leadbetter JR, Newman DK, Readhead C, Bronner ME, Wold B, Lansford R, Sauka-spengler T, Fraser SE, Pierce NA. 2016. Mapping a multiplexed zoo of mRNA expression. Development 143:3632–3637. doi:10.1242/dev.140137

Choi HMT, Schwarzkopf M, Fornace ME, Acharya A, Artavanis G, Stegmaier J, Cunha A, Pierce NA. 2018. Third-generation in situ hybridization chain reaction : multiplexed, quantitative, sensitive, versatile, robust. Development 145:dev165753. doi:10.1242/dev.165753

Chotard C, Salecker I. 2007. Glial cell development and function in the Drosophila visual system. Neuron Glia Biol 3:17–25. doi:10.1017/S1740925X07000592

Csordás G, Grawe F, Uhlirova M. 2020. Eater cooperates with Multiplexin to drive the formation of hematopoietic compartments. Elife 9:e57297. doi:10.7554/eLife.57297

Davies AM. 2003. Regulation of neuronal survival and death by extracellular signals during development. EMBO J 22:2537–2545.

Duckhorn JC, Junker IP, Ding Y, Shirangi TR. 2022. Combined in Situ Hybridization Chain Reaction and Immunostaining to Visualize Gene Expression in Whole-Mount Drosophila Central Nervous Systems JuliaBehavioral Neurogenetics. Springer Nature. doi:10.1007/978-1-0716-2321-3

Edwards TN, Nuschke AC, Nern A, Meinertzhagen I a. 2012. Organization and metamorphosis of glia in the Drosophila visual system. J Comp Neurol 520:2067–2085. doi:10.1002/cne.23071

Fernandes VM, Chen Z, Rossi AM, Zipfel J, Desplan C. 2017. Glia relay differentiation cues to coordinate neuronal development in Drosophila. Science (80-) 357:886–891.

Fischbach KF, Dittrich APM. 1989. The optic lobe of Drosophila melanogaster. I: A. Golgi analysis of wild-type structure. Cell Tissue Res 258:441–475. doi:doi: 10.1007/BF00218858

Franzdóttir SR, Engelen D, Yuva-Aydemir Y, Schmidt I, Aho A, Klämbt C. 2009. Switch in FGF signalling initiates glial differentiation in the Drosophila eye. Nature 460:758–761. doi:10.1038/nature08167

Freeman M, Klämbt C, Goodman CS, Rubin GM. 1992. The argos gene encodes a diffusible factor that regulates cell fate decisions in the drosophila eye. Cell 69:963–975. doi:10.1016/0092-8674(92)90615-J

Fuchs Y, Steller H. 2011. Programmed Cell Death in Animal Development and Disease. Cell 147:742–758. doi:10.1016/j.cell.2011.10.033

Golembo M, Schweitzer R, Freeman M, Shilo B. 1996. argos transcription is induced by the Drosophila EGF receptor pathway to form an inhibitory feedback loop. Development 230:223–230.

Hadjieconomou D, Timofeev K, Salecker I. 2011. A step-by-step guide to visual circuit assembly in Drosophila. Curr Opin Neurobiol 21:76–84. doi:10.1016/j.conb.2010.07.012

Hasegawa E, Kaido M, Takayama R, Sato M. 2013. Brain-specific-homeobox is required for the specification of neuronal types in the Drosophila optic lobe. Dev Biol 377:90–99. doi:10.1016/j.ydbio.2013.02.012

Hennig KM, Colombani J, Neufeld TP. 2006. TOR coordinates bulk and targeted endocytosis in the Drosophila melanogaster fat body to regulate cell growth . J Cell Biol 173:963–974. doi:10.1083/jcb.200511140

Hidalgo A. 2003. The control of cell number during central nervous system development in flies and mice 120:1311–1325. doi:10.1016/j.mod.2003.06.004

Huang Z, Kunes S. 1998. Signals transmitted along retinal axons in Drosophila: Hedgehog signal reception and the cell circuitry of lamina cartridge assembly. Development 125:3753–64. doi:10.1016/s0092-8674(00)80094-x

Huang Z, Kunes S. 1996. Hedgehog, transmitted along retinal axons, triggers neurogenesis in the developing visual centers of the Drosophila brain. Cell 86:411–22. doi:10.1016/S0092-8674(00)80114-2

Huang Z, Shilo BZ, Kunes S. 1998. A retinal axon fascicle uses spitz, an EGF receptor ligand, to construct a synaptic cartridge in the brain of Drosophila. Cell 95:693–703. doi:10.1016/S0092-8674(00)81639-6

Jenett A, Rubin GM, Ngo T-TB, Shepherd D, Murphy C, Dionne H, Pfeiffer BD, Cavallaro A, Hall D, Jeter J, Iyer N, Fetter D, Hausenfluck JH, Peng H, Trautman ET, Svirskas RR, Myers EW, Iwinski ZR, Aso Y, DePasquale GM, Enos A, Hulamm P, Lam SCB, Li H-H, Laverty TR, Long F, Qu L, Murphy SD, Rokicki K, Safford T, Shaw K, Simpson JH, Sowell A, Tae S, Yu Y, Zugates CT. 2012. A GAL4-driver line resource for Drosophila neurobiology. Cell Rep 2:991–1001. doi:10.1016/j.celrep.2012.09.011

Kamimura K, Koyama T, Habuchi H, Ueda R, Masu M, Kimata K, Nakato H. 2006. Specific and flexible roles of heparan sulfate modifications in Drosophila FGF signaling . J Cell Biol 174:773–778. doi:10.1083/jcb.200603129

Kurada P, White K. 1998. Ras Promotes Cell Survival in Drosophila by Downregulating hid Expression 95:319–329.

Louradour I, Sharma A, Morin-Poulard I, Letourneau M, Vincent A, Crozatier M, Vanzo N. 2017. Reactive oxygen species-dependent Toll/NF-κB activation in the Drosophila hematopoietic niche confers resistance to wasp parasitism. Elife 6:e25496. doi:10.7554/eLife.25496

Luo L, Flanagan JG. 2007. Development of Continuous and Discrete Neural Maps. Neuron 56:284–300. doi:10.1016/j.neuron.2007.10.014

Malin J, Desplan C. 2021. Neural specification, targeting, and circuit formation during visual system assembly. Proc Natl Acad Sci 118:e21018231182. doi:10.1073/pnas.2101823118

Miguel-Aliaga I, Thor S. 2009. Programmed cell death in the nervous system — a programmed cell fate ? Curr Opin Neurobiol 19:127–133. doi:10.1016/j.conb.2009.04.002

Morante J, Vallejo DM, Desplan C, Dominguez M. 2013. Conserved miR-8/miR-200 Defines a Glial Niche that Controls Neuroepithelial Expansion and Neuroblast Transition. Dev Cell 27:174–187. doi:10.1016/j.devcel.2013.09.018

Park H, Poo MM. 2013. Neurotrophin regulation of neural circuit development and function. Nat Rev Neurosci 14:7–23. doi:10.1038/nrn3379

Pastor-Pareja JC, Xu T. 2011. Shaping Cells and Organs in *Drosophila* by Opposing Roles of Fat Body-Secreted Collagen IV and Perlecan. Dev Cell 21:245–256. doi:10.1016/j.devcel.2011.06.026

Pinto-Teixeira F, Konstantinides N, Desplan C. 2016. Programmed cell death acts at different stages of Drosophila neurodevelopment to shape the central nervous system. FEBS Lett 590:2435–2453. doi:10.1002/1873-3468.12298.Programmed

Ren Q, Awasaki T, Wang Y, Huang Y, Lee T. 2018. Lineage-guided Notch-dependent gliogenesis by Drosophila multi-potent progenitors. Development 145:dev160127. doi:10.1242/dev.160127

Schindelin J, Arganda-carreras I, Frise E, Kaynig V, Longair M, Pietzsch T, Preibisch S, Rueden C, Saalfeld S, Schmid B, Tinevez J, White DJ, Hartenstein V, Eliceiri K, Tomancak P, Cardona A. 2012. Fiji : an open-source platform for biological-image analysis. Nat Methods 9:676–682. doi:10.1038/nmeth.2019

Sopko R, Perrimon N. 2013. Receptor tyrosine kinases in Drosophila development. Cold Spring Harb Perspect Biol 5:a009050.

Spéder P, Brand AH. 2014. Gap junction proteins in the blood-brain barrier control nutrient- dependent reactivation of Drosophila neural stem cells. Dev Cell 30:309–321. doi:10.1016/j.devcel.2014.05.021

Sugie A, Umetsu D, Yasugi T, Fischbach KF, Tabata T. 2010. Recognition of pre- and postsynaptic neurons via nephrin/NEPH1 homologs is a basis for the formation of the Drosophila retinotopic map. Development 137:3303–3313. doi:10.1242/dev.047332

Tamamouna V, Rahman MM, Petersson M, Charalambous I, Kux K, Mainor H, Bolender V, Isbilir B, Edgar BA, Pitsouli C. 2021. Remodelling of oxygen-transporting tracheoles drives intestinal regeneration and tumorigenesis in Drosophila. Nat Cell Biol 23:497–510. doi:10.1038/s41556-021-00674-1

Tan L, Zhang KX, Pecot MY, Nagarkar-Jaiswal S, Lee PT, Takemura SY, McEwen JM, Nern A, Xu S, Tadros W, Chen Z, Zinn K, Bellen HJ, Morey M, Zipursky SL. 2015. Ig Superfamily Ligand and Receptor Pairs Expressed in Synaptic Partners in Drosophila. Cell 163:1756–1769. doi:10.1016/j.cell.2015.11.021

Treisman JE. 2013. Retinal differentiation in Drosophila. Wiley Interdiscip Rev Dev Biol 2:545–557. doi:10.1002/wdev.100

Tsruya R, Schlesinger A, Reich A, Gabay L, Sapir A, Shilo B. 2002. Intracellular trafficking by Star regulates cleavage of the Drosophila EGF receptor ligand Spitz. Genes Dev 12:222–234. doi:10.1101/gad.214202.fate

Umetsu D, Murakami S, Sato M, Tabata T. 2006. The highly ordered assembly of retinal axons and their synaptic partners is regulated by Hedgehog/Single-minded in the Drosophila visual system. Development 133:791–800. doi:10.1242/dev.02253

Urban S, Lee JR, Freeman M. 2002. A family of rhomboid intramembrane proteases activates all Drosophila membrane-tethered EGF ligands. EMBO J 21:4277–4286. doi:10.1093/emboj/cdf434

Viktorin G, Riebli N, Reichert H. 2013. A multipotent transit-amplifying neuroblast lineage in the central brain gives rise to optic lobe glial cells in Drosophila. Dev Biol 379:182–194. doi:10.1016/j.ydbio.2013.04.020

Wu B, Li J, Chou Y, Luginbuhl D, Luo L. 2017. Fibroblast growth factor signaling instructs ensheathing glia wrapping of Drosophila olfactory glomeruli. Proc Natl Acad Sci 114:7505–7512. doi:10.1073/pnas.1706533114

Yamaguchi Y, Miura M. 2015. Review Programmed Cell Death in Neurodevelopment. Dev Cell 32:478–490. doi:10.1016/j.devcel.2015.01.019

Yogev S, Schejter ED, Shilo B-Z. 2008. Drosophila EGFR signalling is modulated by differential compartmentalization of Rhomboid intramembrane proteases. EMBO J 27:1219–1230. doi:https://doi.org/10.1038/emboj.2008.58

Yogev S, Schejter ED, Shilo BZ. 2010. Polarized secretion of drosophila EGFR ligand from photoreceptor neurons is controlled by ER localization of the ligand-processing machinery. PLoS Biol 8:e1000505.

Zang Y, Wan M, Liu M, Ke H, Ma S, Liu L, Pastor-pareja C. 2015. Plasma membrane overgrowth causes fibrotic collagen accumulation and immune activation in Drosophila adipocytes. Elife 4:e07187. doi:10.7554/eLife.07187

