## Supplementary File 1 for "Differentiation signals from glia are fine-tuned to set neuronal numbers during development"

**Supplementary File 1: Table 1 summarising the results from the glial-Gal4 screen (Figure 1 B,C, Figure 1- figure supplement 1B-N) to identify the glial type that regulates L5 development.**

| ***Glial-type > EGFR^DN^x2*** | **Elav+ cells in the proximal lamina** |
| --- | --- |
| Pan-glia | Absent or reduced |
| Perineurial glia | Present |
| Sub-perineurial glia | Present |
| Cortex glia | Present |
| Epithelial glia and Marginal glia (eg + mg) | Present |
| Wrapping glia and xg^outer^ (xg^O^) | Absent or reduced |
| xg^O^ +xg^inner^ | Absent or reduced |
| xg^O^ | Absent or reduced |
| xg^Ots^ | Absent or reduced |
