## Supplementary File 2 for "Differentiation signals from glia are fine-tuned to set neuronal numbers during development"

**Supplementary File 2: Table listing all genotypes and experimental conditions used by figure panel.** (Note that only female genotypes are listed though both sexes were included in our analyses)

| **Figure** | **Panel** | **Genotype** | **Conditions** |
| --- | --- | --- | --- |
| 1 | B | *y,w/+ ;UAS-CD8::GFP/+; hh-Gal4/repo-QF2,10xQUAS-6xm.Cherry* | Raised at 25^o^C |
| 1 | C | *w^1118^/+; tub-Gal80ts/+; Repo-Gal4/UAS-CD8::GFP* | Raised at 18^o^C till late-L2, then shifted to 29^o^C for 48 hours |
| 1 | D | *y,w/+; UAS-CD8::GFP/ UAS-EGFR^DN^; Repo-Gal4/ UAS-EGFR^DN^* | Raised at 18^o^C till late-L2, then shifted to 29^o^C for 48 hours |
| 1 | E | *y,w/+; UAS-CD8::GFP/+; R25A01-Gal4/+* | Raised at 29^o^C |
| 1 | F | *y,w/+; UAS-CD8::GFP/ UAS-EGFR^DN^; R25A01-Gal4/ UAS-EGFR^DN^* | Raised at 29^o^C |
| 1 | G | *w^1118^/+;; R25A01-Gal4/UAS-lacZ* | Raised at 29^o^C |
| 1 | H | *w^1118^/+; UAS-EGFR^DN^/+; R25A01-Gal4/ UAS-EGFR^DN^* | Raised at 29^o^C |
| 1- fig. supp.1 | B | *y,w/+; NP6293-Gal4/+; 10xUAS-myrGFP/+* | Raised at 29^o^C |
| 1- fig. supp.1 | C | *y,w/+; NP6293-Gal4/UAS-EGFR^DN^; UAS-EGFR^DN^/+* | Raised at 29^o^C |
| 1- fig. supp.1 | D | *w/+; NP2276-Gal4/+; 10xUAS-myrGFP/+* | Raised at 29^o^C |
| 1- fig. supp.1 | E | *w/+; NP2276-Gal4/ UAS-EGFR^DN^; UAS-EGFR^DN^/+* | Raised at 29^o^C |
| 1- fig. supp.1 | F | *w^1118^/+;; R54H02-Gal4/10xUAS-myrGFP* | Raised at 29^o^C |
| 1- fig. supp.1 | G | *w^1118^/+; UAS-EGFR^DN^/+; R54H02-Gal4/ UAS-EGFR^DN^* | Raised at 29^o^C |
| 1- fig. supp.1 | H | *w^1118^/+;; R10C12-Gal4/10xUAS-myrGFP* | Raised at 29^o^C |
| 1- fig. supp.1 | I | *w^1118/+^; UAS-EGFR^DN^/+; R10C12-Gal4/UAS-EGFR^DN^* | Raised at 29^o^C |
| 1- fig. supp.1 | J | *w/+; Mz97-Gal4, UAS-Stinger/+; 10xUAS-myrGFP/+* | Raised at 29^o^C |
| 1- fig. supp.1 | K | *w/+; Mz97-Gal4, UAS-Stinger/UAS-EGFR^DN^; UAS-EGFR^DN^/+* | Raised at 29^o^C |
| 1- fig. supp.1 | L | *w^1118^/+;; R53H12-Gal4/10xUAS-myrGFP* | Raised at 29^o^C |
| 1- fig. supp.1 | M | *w^1118^/+; UAS-EGFR^DN^/+; R53H12-Gal4/ UAS-EGFR^DN^* | Raised at 29^o^C |
| 1- fig. supp.1 | N | *y,w/+; tub-Gal80^ts^/+; R25A01-Gal4/UAS-EGFR^DN^* | Raised at 29^o^C |
| 1- fig. supp.1 | O,Q | *y,w/+; UAS-LifeAct-GFP/UAS-nls.lacZ; R25A01-Gal4/UAS-nls.lacZ* | Raised at 29^o^C |
| 1- fig. supp.1 | P,R | *y,w,hsflp^122^/+; UAS-LifeAct-GFP/UAS-EGFR^DN^; R25A01-Gal4/UAS-EGFR^DN^* | Raised at 29^o^C |
| 1- fig. supp.1 | S | *Canton S* | Raised at 29^o^C |
| 1- fig. supp.1 | T | *y,w/+; UAS-EGFR^DN^/+; R25A01-Gal4/UAS-EGFR^DN^* | Raised at 29^o^C |
| 2 | A | *w^1118^/+;; R25A01-Gal4/UAS-lacZ* | Raised at 29^o^C |
| 2 | B,C,D | *w^1118^/+;; R25A01-Gal4/ UAS-EGFR^DN^* | Raised at 29^o^C |
| 3 | A | *y,w; GMR-Gal4/+; rho3^PLLb^, UAS-CD8::GFP* | Raised at 29^o^C |
| 3 | B,C | *y,w; GMR-Gal4/UAS-rho3-3xHA; rho3^PLLb^, UAS-CD8::GFP* | Raised at 29^o^C |
| 3 | D,E | *w^1118^/+;; R25A01-Gal4/UAS-CD8::GFP* | Raised at 29^o^C |
| 3 | F,G | *w^1118^/+; UAS-EGFR^DN^/+; R25A01-Gal4/ UAS-CD8::GFP* | Raised at 29^o^C |
| 3 | H,I | *w^1118^/+; UAS-EGFR^DN^/+; R25A01-Gal4/ UAS-m.spi* | Raised at 29^o^C |
| 3 | J,K | *w^1118^/+; UAS-EGFR^DN^/+; R25A01-Gal4/UAS-Cg25c::RFP* | Raised at 29^o^C |
| 3 | L,M | *w^1118^/+;UAS-EGFR^DN^/UAS-nls.lacZ; R25A01-Gal4/UAS-nls.lacZ* | Raised at 29^o^C |
| 3 | N,O | *w^1118^/+;UAS-EGFR^DN^/ UAS-Col4a1^EY11094^; R25A01-Gal4/UAS-m.spi* | Raised at 29^o^C |
| 3 | Q | *w^1118^/+;; R25A01-Gal4/UAS-m.spi* | Raised at 29^o^C |
| 3 | R | *w^1118^/+;; R25A01-Gal4/UAS-Cg25c::RFP* | Raised at 29^o^C |
| 3 | T | *w^1118^/+;UAS-Dcr2/UAS-SpiRNAi; R25A01-Gal4/+* | Raised at 29^o^C |
| 3 | U | *w^1118^/+;UAS-Dcr2/UAS-Col4a1RNAi^v28369^; R25A01-Gal4/+* | Raised at 29^o^C |
| 3 | V | *w^1118^/+;UAS-Dcr2/UAS-SpiRNAi; R25A01-Gal4/UAS-Col4a1RNAi^v104536^* | Raised at 29^o^C |
| 3-fig. supp. 1 | A | *w^1118^/+;; R25A01-Gal4/UAS-lacZ* | Raised at 29^o^C |
| 3-fig. supp. 1 | B | *y,w/+; UAS-EGFR^DN^/+; R25A01-Gal4/UAS-nls.lacZ* | Raised at 29^o^C |
| 3-fig. supp. 1 | C | See list of *Drosophila* stocks for specific strains used. | Raised at 29^o^C |
| 3-fig. supp. 1 | D | *w/+;; bnl^NP2211^-Gal4/ UAS-CD8::GFP* | Raised at 29^o^C |
| 3-fig. supp. 1 | E | *w/+; ths^MI07139^-Gal4/+; MKRS/UAS-CD8::GFP* | Raised at 29^o^C |
| 3-fig. supp. 1 | F | *y,w/+; Cg-Gal4/+; UAS-CD8::GFP/+* | Raised at 29^o^C |
| 3-fig. supp. 1 | G | *y,w/+; Spi^NP0289^-Gal4/+; UAS-CD8::GFP/+* | Raised at 29^o^C |
| 3-fig. supp. 1 | H, I | *w^1118^/+;; R25A01-Gal4/UAS-CD8::GFP* | Raised at 29^o^C |
| 3-fig. supp. 1 | J | *w^1118^/+; UAS-EGFR^DN^/+; R25A01-Gal4/ UAS-s.spi* | Raised at 29^o^C |
| 3-fig. supp. 1 | L,N | *y,w/+; UAS-CD8::GFP/ UAS-EGFR^DN^; R25A01-Gal4/ UAS-EGFR^DN^* | Raised at 29^o^C |
| 3-fig. supp. 1 | P | *w^1118^/+; Ddr^CR01018^-Gal4/+; UAS-lacZ/+* | Raised at 29^o^C |
| 3-fig. supp. 1 | Q | *Canton S* | Raised at 25^o^C |
| 3-fig. supp. 1 | R | *w^1118^/+; UAS-EGFR^DN^/+; R25A01-Gal4/ UAS-m.spi* | Raised at 29^o^C |
| 3-fig. supp. 1 | S | *w^1118^/+; UAS-EGFR^DN^/+; R25A01-Gal4/UAS-Cg25c::RFP* | Raised at 29^o^C |
| 3- fig. supp. 2 | A | *w^1118^/+;; R25A01-Gal4/UAS-lacZ* | Raised at 29^o^C |
| 3-fig. supp. 2 | B | *w^1118^/+;UAS-Dcr2/UAS-SpiRNAi; R25A01-Gal4/UAS-Col4a1RNAi^v104536^* | Raised at 29^o^C |
| 4 | A,B | *Canton S* | Raised at 25^o^C |
| 4 | C,D | *ey-Gal80/+; tub-Gal80^ts^/+; R27G05/UAS-PntP1* | Raised at 18^o^C till the late second larval instar (L2), then shifted to 29^o^C for 48 hours |
| 4-fig. supp. 1 | A,B | *ey-Gal80/+; tub-Gal80^ts^/+; R27G05/UAS-lacZ* | Raised at 18^o^C till the late second larval instar (L2), then shifted to 29^o^C for 48 hours |
| 4-fig. supp. 1 | C,D | *ey-Gal80/+; tub-Gal80^ts^/UAS-rl^SEM^; R27G05/+* | Raised at 18^o^C till the late second larval instar (L2), then shifted to 29^o^C for 48 hours |
| 5 | A,B | *w;;FRT80B, Dronc^I24^* | Raised at 25^o^C |
| 5 | D,E | *;;aos^w11^/+* | Raised at 29^o^C |
| 5 | F | *w^1118^; R64B07-Gal4/UAS-nls.lacZ; UAS-Dcr-2/+* | Raised at 29^o^C |
| 5 | G | *w^1118^; R64B07-Gal4/UAS-aos^RNAi^; UAS-Dcr-2/+* | Raised at 29^o^C |
| 5- fig. supp.1 | A | *w^1118^/+;R64B07-Gal4/+;UAS-CD8::GFP/+* | Raised at 29^o^C |
